# Heterogeneity-Resolved Ultrafast Transient Absorption Spectroscopy of Single Supramolecular Light-Harvesting Antennas

**DOI:** 10.64898/2026.01.19.700341

**Authors:** Shun Arai, Shogo Matsubara, Toru Kondo

**Affiliations:** Division of Photophysical Biology, National Institute for Basic Biology, National Institutes of Natural Sciences, Okazaki, Aichi 444-8787, Japan; The Graduate University for Advanced Studies, SOKENDAI, Okazaki, Aichi 444-8787, Japan; Department of Life Science and Applied Chemistry, Graduate School of Engineering, Nagoya Institute of Technology, Nagoya, Aichi 466-8555, Japan; Interconnective Photobiology Group, Exploratory Research Center on Life and Living Systems, National Institutes of Natural Sciences, Okazaki, Aichi 444-8787, Japan

## Abstract

Photosynthetic light harvesting proceeds rapidly and robustly through precisely arranged pigment molecules. However, the molecular arrangements are not uniform because they exhibit structural heterogeneity among individual complexes and across spatial regions, and undergo dynamic fluctuations. Such static and dynamic disorder can substantially perturb excitation dynamics. Here, we report a highly sensitive transient absorption microscope that integrates single-objective absorption microscopy, balanced detection, and lock-in amplification, enabling quantitative analysis of heterogeneity and temporal fluctuations in excitation dynamics. By analyzing individual chlorophyll-derivative aggregates mimicking photosynthetic light-harvesting antennas, we demonstrated that two kinetic components with nearly identical time constants can be resolved based on differences in their time-constant distributions. We further quantified the photophysical properties of each component, including absorbance change, fluorescence intensity, fluorescence efficiency, and fluorescence peak intensity ratio. These results establish an analytical framework for excitation dynamics that leverages not only the mean values of time constants but also their distribution profiles.

## MAIN TEXT

Photosynthetic organisms have evolved light-harvesting antenna complexes that functionally organize pigment molecules within protein scaffolds to efficiently capture light energy. Advances in structural biology and spectroscopy have elucidated pigment–protein and pigment–pigment interactions, thereby deepening our understanding of their structure–function relationships. However, protein scaffolds undergo continuous thermal fluctuations, giving rise to dynamic structural disorder.^1-2^ In addition, static disorder manifests as structural heterogeneity, such that even assemblies composed of identical protein components can adopt distinct structures.^3-4^ Furthermore, chlorosomes, the light-harvesting antennas of green sulfur bacteria, are known to consist of protein-free pigment aggregates that exhibit pronounced structural heterogeneity.^5-7^ Despite this structural disorder, photosynthetic light-harvesting antennas maintain remarkably high efficiency and functional robustness, yet the underlying molecular mechanisms remain elusive.

Addressing this issue requires the experimental quantification of the respective contributions of dynamic and static structural disorder to excitation dynamics. However, such subtle differences are obscured by ensemble averaging in conventional measurements. To overcome this limitation, single-molecule spectroscopy (SMS) has emerged. In particular, fluorescence-based SMS is widely utilized in biological research,^8-9^ because it achieves nearly background-free, high-sensitivity detection. Numerous applications to photosynthetic light-harvesting complexes have elucidated spectral variations arising from structural heterogeneity,^6,10-11^ revealed photoprotection via non-photochemical quenching (NPQ),^12-15^ and identified the lowest-energy chlorophylls (Chls) in light-harvesting systems.^16-17^

Meanwhile, fluorescence-based SMS faces challenges in resolving ultrafast processes. Consequently, femtosecond time-resolved SMS has been demonstrated by inducing stimulated emission using a pair of pulses and tracking the excited-state populations via fluorescence intensity.^18-19^ This approach enables the observation of ultrafast energy transfer and relaxation processes in individual photosynthetic antenna complexes.^20-22^ Nevertheless, it is difficult to analyze multistep energy transfer processes within complex pigment architectures and to quantify the parameters governing excitation dynamics. Furthermore, non-fluorescent molecules and dark states are inaccessible as long as measurements rely exclusively on fluorescence detection, which presents a fundamental limitation.

Transient absorption (TA) spectroscopy, a complementary approach to fluorescence spectroscopy, is suitable for observing excitation energy transfer, exciton dynamics, and radical intermediates formed during charge separation and electron transfer.^23-26^ Combining TA with microscopy has enabled spatiotemporally-resolved measurements.^27-31^ Additionally, single-molecule steady-state absorption has been observed.^32-34^ However, extending TA to the single-molecule level remains highly challenging, particularly for biological and organic samples susceptible to photobleaching, because it requires ultrahigh sensitivity together with high spatial and temporal resolution. Moreover, capturing temporal variations in excitation dynamics associated with structural fluctuations demands continuous acquisition of TA time-series data. Here, we report a TA microscope with high sensitivity that approaches the single-molecule level, as well as diffraction-limited spatial resolution and femtosecond temporal resolution. Using this instrument, we performed TA measurements on individual supramolecular light-harvesting antennas that mimic the pigment aggregates in chlorosomes. The observed kinetic heterogeneity enables quantitative resolution and characterization of multiple kinetic components attributable to structural disorder in excitation dynamics.

**Figure 1** shows a schematic diagram of the developed TA microscope. Pulses emitted from a Ti:sapphire laser (Synergy, Spectra Physics), with a pulse duration of 14 fs and a repetition rate of 78 MHz, were separated into pump and probe beams by a beam splitter. Their wavelengths were selected using band-pass filters centered at 720 nm (FF01-720/24-25, Semrock) and 750 nm (FBH750-10, Thorlabs), respectively. The degrees of polarization of the pump and probe light were reduced to 0.09 and 0.25, respectively, by quarter-wave plates (WPM03-Q-NIR, Newlight Photonics) (see **Supporting Information** Section 1). The pump light was modulated at 45 kHz by an optical chopper (310CD, Scitec Instruments) after being reflected by a dichroic mirror (FELH0750, Thorlabs). In the probe path, a rapid delay stage (scanDelay USB-15, APE) was introduced to scan the delay time between the pump and probe pulses at 4 Hz over a range of 14.4 ps, providing an effective measurement time window of 12 ps. These pump and probe beams were spatially overlapped at a beam splitter and subsequently separated by another beam splitter into two beams, i.e., reference and signal beams. The pump light in the reference beam was removed by long-pass filters (FELH0750, Thorlabs). Thus, only the probe light was detected by the reference channel of a balanced photodiode (Nirvana, Newport). In the signal arm, the signal beam was focused onto the sample placed on a piezo stage (Nano-LP100, Mad City Labs) using an oil-immersion objective with an NA = 1.45 (UPLXAPO100XO, Olympus), where the beam exiting the objective was reflected by a mirror positioned above the sample and subsequently focused onto the sample (**Figure 1**, inset). This optical configuration differs from that of conventional absorption microscopes, which employ two objectives arranged face to face, with one focusing the beam onto the sample and the other collecting the transmitted beam.^33,35^ In our TA microscope, a single objective both focuses and collects the beam, eliminating instability in the relative position between the two objectives. The signal beam transmitted through the sample propagated in the reverse direction and passed through the beam splitter. As in the reference arm, the pump light in the signal beam was removed by long-pass filters (FELH0750, Thorlabs), and only the probe light was detected by the signal channel of the balanced photodiode.

**Figure 1.**
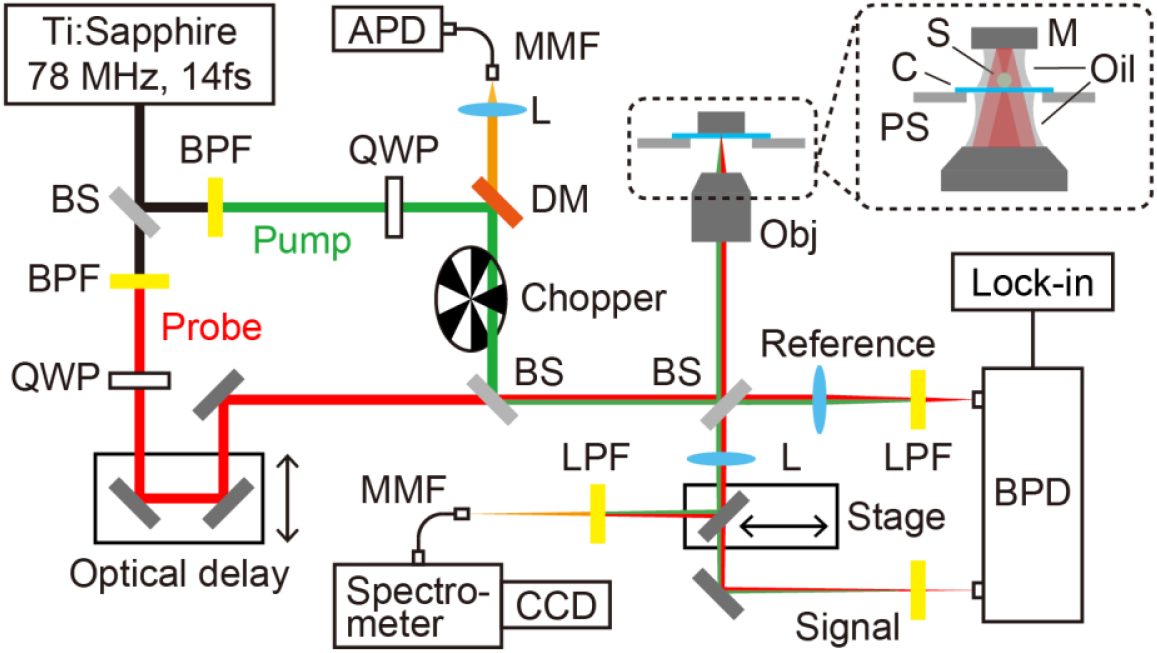
Schematic of the developed TA microscope. BS: beam splitter; BPF: band-pass filter; QWP: quarter-wave plate; DM: dichroic mirror; L: lens; Obj: objective; S: sample; C: coverslip; Oil: immersion oil; PS: piezo stage; M: mirror; LPF: long-pass filter; MMF: multimode fiber; APD: avalanche photodiode; BPD: balanced photodiode; CCD: charge-coupled device; Lock-in: lock-in amplifier. The double-headed arrows indicate translations of the mirror-mounted stage used for scanning the pump–probe delay and for redirecting fluorescence. The inset shows an enlarged view around the sample, where the incident beam is focused onto the sample after being reflected by the mirror, transmitted through the sample, and subsequently collected by the objective.

A lock-in amplifier (LI5640, NF Corporation) demodulated and amplified the signal from the balanced photodiode, which was modulated at the chopping frequency. The digitizer board (DIG-100M1002-PCI, CONTEC) acquired data at a sampling rate of 6 kHz, corresponding to a sampling interval of 9.6 fs in pump–probe delay. The TA signals were obtained by averaging data points over a 30 fs interval. The sensitivity limit, defined as the absorbance change yielding a signal-to-noise ratio (SNR) of 1, was estimated to be ∼10^−7^ at a probe power of 81 µW and a lock-in time constant of 30 ms (**Figure S1**). Given the absorbance of the photosynthetic pigment Chl *a*, calculated to be 2.11×10^−7^ from its molar extinction coefficient of 8.13×10^4^ M^−1^ cm^−1^ at the Q_y_ band,^36^ the sensitivity of this instrument approaches the single-molecule level (see **Supporting Information** Section 2). In the present measurements, the incident power was reduced to 14 µW to avoid photodamage, and the lock-in time constant was shortened to 10 µs to enable fast sampling during the rapid delay scan. The sensitivity under this condition was on the order of 10^−4^ (**Figure S1**), which was sufficient for this study.

Fluorescence emitted from the sample was reflected by the mirror positioned above the sample and then collimated by the objective. After propagating back along the same path, it was split into reflected and transmitted beams by the beam splitter. The reflected fluorescence passed through the dichroic mirror and was detected by an avalanche photodiode (SPCM-AQRH-14-FC, Excelitas) to acquire fluorescence images. The transmitted fluorescence was redirected by translating the mirror and recorded with a charge-coupled device (CCD) spectrometer (Kymera328i and iDus416, Andor) to acquire fluorescence spectra. The excitation wavelength was set to ∼695 nm using a short-pass filter (FESH0700, Thorlabs). The spatial resolution, defined as the full width at half maximum (FWHM) of the focal spot, was estimated to be 336, 243, and 1145 nm for absorption and 309, 257, and 1031 nm for fluorescence along the *X*-, *Y*-, and *Z*-axes, respectively (see **Supporting Information** Section 3).

Using the developed setup, we analyzed supramolecular light-harvesting antennas that mimic the bacteriochlorophyll (BChl) self-aggregates in chlorosomes. A Chl derivative, Zn-HM, in which the central Mg is replaced by Zn (**Figure 2a**), self-assembles into amorphous particles (**Figure 2b**) (see **Supporting Information** Section 4).^37^ When the solution is left undisturbed for more than a week, the assemblies become thermodynamically stabilized, forming well-ordered, unidirectionally stacked tubes (**Figure 2c**). The particle-like and tubular aggregates have both been observed by atomic force microscopy (AFM).^37^ Thus, the Zn-HM self-aggregates are likely composed of a heterogeneous mixture of highly ordered and less ordered structures, raising the question of how this local structural heterogeneity contributes to excitation dynamics. To address this, TA measurements were performed in local regions within individual aggregates.

**Figure 2.**
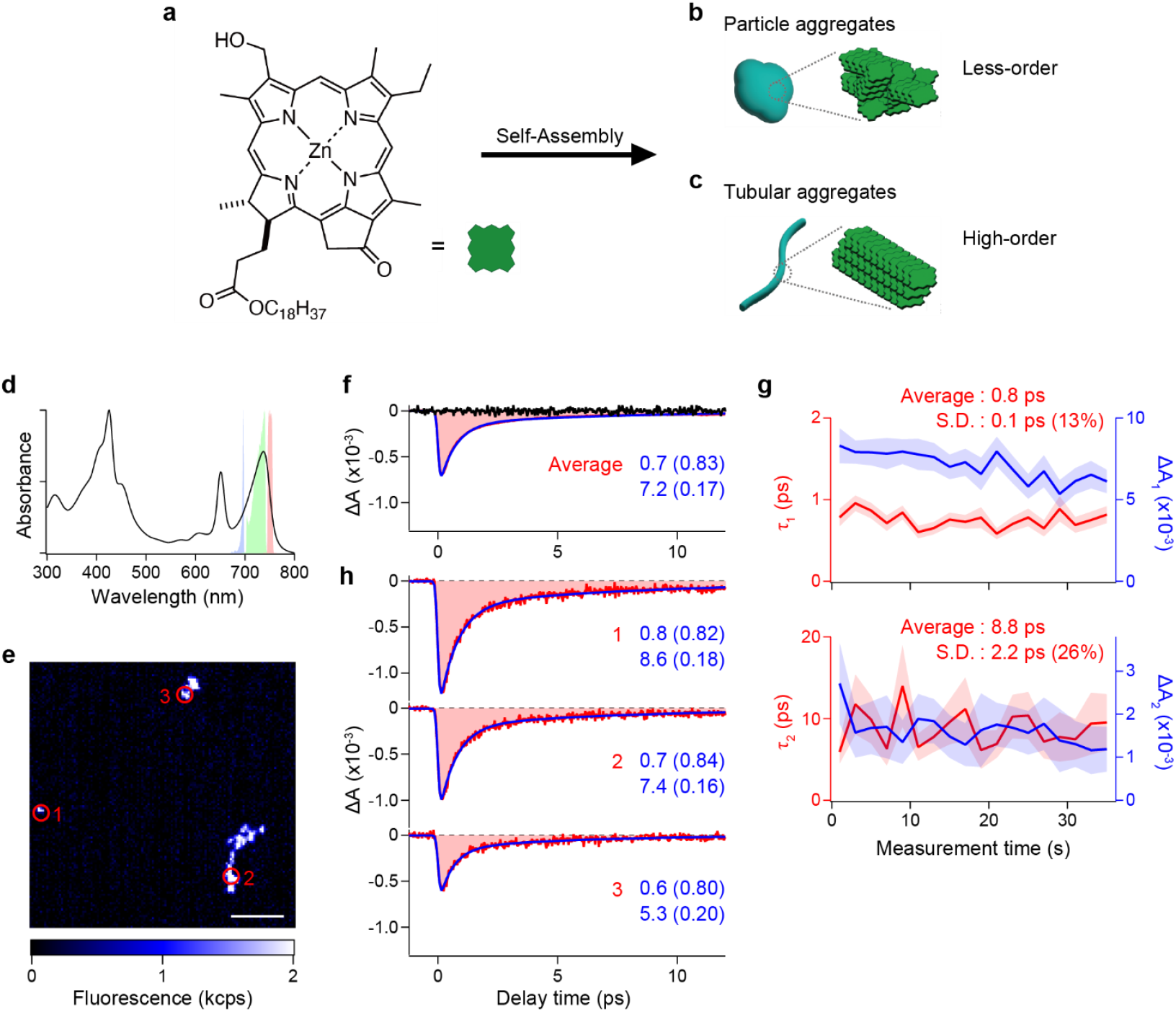
(a–c) Molecular structure of Zn-HM (a) and schematic illustrations of self-assembled particle-like (b) and tubular (c) aggregates. (d) Absorption spectrum of the Zn-HM aggregates, together with spectra of the excitation light (blue) for fluorescence imaging and spectral measurements and of the pump (green) and probe (red) light for TA measurements. (e) Fluorescence image of the Zn-HM aggregates. The scale bar represents 5 µm. (f) Ensemble-averaged TA signal (red). The background signal (black) and the fitted curve (blue) are overlaid. (g) Temporal variations in *τ*_1_ and Δ*A*_1_ (top panel, red and blue, respectively) and *τ*_2_ and Δ*A*_2_ (bottom panel, red and blue, respectively) measured at site 1 in (e). Mean values, S.D., and coefficients of variation for *τ*_1_ and *τ*_2_ are indicated in each panel. The shaded areas represent ± one standard error for the fit at each time point. (h) TA signals (red) and fitted curves (blue) measured at distinct sites 1–3 in (e). Estimated *τ*_1_ and *τ*_2_ values, with their relative amplitudes in parentheses, are indicated in each panel.

**Figure 2d** shows the absorption spectrum of the Zn-HM aggregates. Whereas the narrow peak at 651 nm is attributed to Zn-HM monomers, the red-shifted peak at 735 nm indicates the formation of J-aggregates.^37^ The aggregates were fixed onto a coverslip by drop casting, followed by spin-coating with a 40 mM Tris-HCl (pH 8) solution containing 1% polyvinyl alcohol (PVA) and an oxygen-scavenging system. Moreover, an Ag mirror coated with immersion oil was placed onto the sample (**Figure 1**, inset, and see also **Supporting Information** Section 4). Fluorescence images, shown in **Figure 2e**, were acquired by collecting emission at >750 nm under excitation at ∼695 nm (**Figure 2d**, blue) with an excitation fluence of 4.0×10^13^ photons pulse^−1^ cm^−2^. Consistent with AFM observations,^37^ particle-like and one-dimensional aggregates were observed (**Figure 2e**, sites 1–3).

TA measurements were performed at 145 fluorescent sites. The pump and probe wavelengths were set to 719–741 nm and 746–755 nm, respectively, to monitor the excitation dynamics within the 735 nm absorption peak (**Figure 2d**, green and red, respectively). The pump and probe fluences were 4.4×10^14^ and 1.1×10^15^ photons pulse^−1^ cm^−2^, respectively. **Figure 2f** shows an ensemble-averaged TA signal (red), which was well fitted with two exponential decay components (blue) (see **Supporting Information** Section 5). The fit provided the time constants (*τ*_1_ and *τ*_2_) and amplitudes (Δ*A*_1_ and Δ*A*_2_) of the two decay components, as well as the total amplitude (total Δ*A*), defined as the sum of Δ*A*_1_ and Δ*A*_2_. The instrumental response function (IRF), which defines the temporal resolution, was estimated to be 177 fs from the fit (see **Supporting Information** Section 5).

The TA traces acquired at 0.25-s intervals were averaged into 2-s bins, and each binned trace was fitted to quantify temporal variations in *τ* and Δ*A* (see **Supporting Information** Section 6). At site 1 in **Figure 2e**, *τ*_1_ had a mean of 0.8 ps, a standard deviation (S.D.) of 0.1 ps, and a coefficient of variation (S.D./mean) of 13% (**Figure 2g**, top panel, red). For *τ*_2_, the corresponding values were 8.8 ps, 2.2 ps, and 26%, respectively (**Figure 2g**, bottom panel, red). These temporal variations in excitation kinetics likely reflect contributions from local structural fluctuations, i.e., dynamic disorder, within the aggregates. Additionally, Δ*A* fluctuated and gradually decreased due to photobleaching (**Figure 2g**, blue). At sites 2 and 3 in **Figure 2e**, the coefficient of variation for *τ*_1_ (*τ*_2_) was estimated to be 19% (43%) and 28% (52%), respectively, which differed from those at site 1 (see **Supporting Information** Section 6). These variations across sites and aggregates suggest that the extent of dynamic disorder differs among sites and aggregates.

At each measurement site, all TA traces in the time series were averaged and then fitted to a biexponential function. **Figure 2h** shows the results at sites 1–3 in **Figure 2e**. Both the time constants and amplitudes varied across sites, indicating heterogeneity in excitation dynamics arising from subtle local structural differences within the aggregates, i.e., static disorder. The distributions of *τ*_1_ and *τ*_2_ were obtained by fitting data acquired at 143 distinct sites. 14% (20/143) of the *τ*_2_ values exceeded the 12-ps measurement time window. Because estimates of time constants beyond this window are uncertain (see **Supporting Information** Section 7), statistical analyses involving *τ*_2_ were restricted to *τ*_2_ < 12 ps. The histogram of *τ*_1_ exhibited a peak at 0.7 ps with broad tails on both sides (**Figure 3a**), implying that the distribution contains both narrow and broad components. Similarly, the histogram of *τ*_2_ appeared to comprise two overlapping components centered around 7.5 ps (**Figure 3b**). Both histograms were well reproduced with two Gaussian components, indicating a mixture of two *τ* distributions with distinct widths (see **Supporting Information** Section 8).

**Figure 3.**
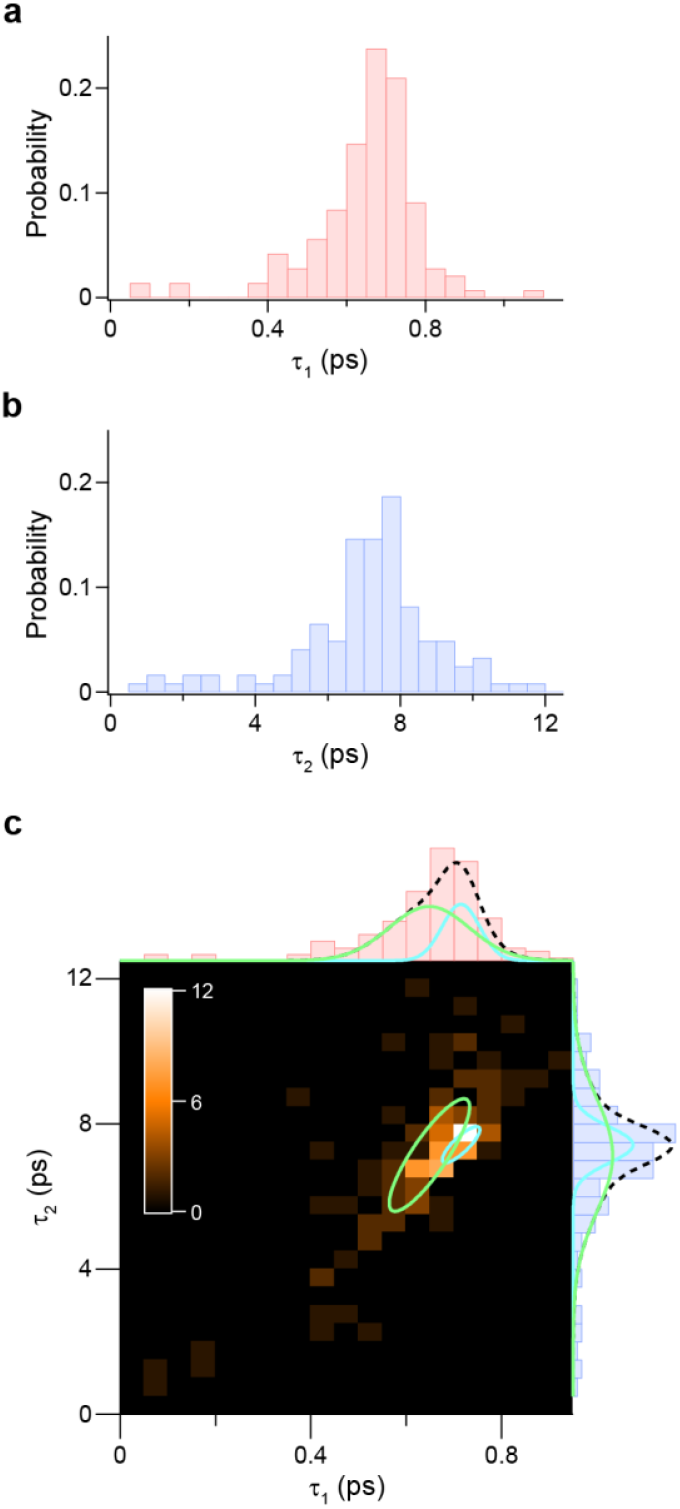
(a, b) Histograms of *τ*_1_ (a) and *τ*_2_ (b), constructed with bin sizes of 0.05 and 0.5 ps, respectively. (c) 2D histogram of *τ*_1_ versus *τ*_2_. Circles indicate the contour at 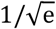 of the peak amplitude of the C_B_ (green) and C_N_ (cyan) components, identified by fitting with a two-component 2D Gaussian function. Projections of the distributions of each component onto the *τ*_1_- and *τ*_2_-axes are shown at the top and right of the panel, respectively. The dashed black lines indicate the summed distributions along each axis.

The relationship between *τ*_1_ and *τ*_2_ was examined by constructing a two-dimensional (2D) histogram of *τ*_1_ versus *τ*_2_ across measurement sites (**Figure 3c**). The pronounced distribution along the diagonal indicates a strong positive correlation between *τ*_1_ and *τ*_2_, suggesting that the two components in their distributions are associated with one another. Therefore, the 2D distribution was fitted with a two-component 2D Gaussian function to identify the two components (see **Supporting Information** Section 8). **Figure 3c** shows two resolved components with broad (green) and narrow (cyan) widths, hereafter denoted as C_B_ and C_N_, respectively. For the C_B_ distribution component, the centers along the *τ*_1_- and *τ*_2_-axes were 0.6 and 7.1 ps, with FWHMs of 0.2 and 3.7 ps, respectively, and the correlation coefficient (*r*) between *τ*_1_ and *τ*_2_ was 0.86. For C_N_, the centers along the *τ*_1_- and *τ*_2_-axes were 0.7 and 7.4 ps, with FWHMs of 0.1 and 1.2 ps, respectively, and *r* was 0.83. The C_B_ population was 67%, which is 2.1 times that of C_N_ (33%). The high *r* values for both C_B_ and C_N_ reflect strong positive correlations between *τ*_1_ and *τ*_2_. While the central *τ* values were nearly identical for C_B_ and C_N_, the FWHMs of *τ*_1_ and *τ*_2_ for C_B_ were 2.1 and 3.1 times as large as those for C_N_, respectively. This demonstrates that even when multiple kinetic components exhibit nearly identical *τ* values, they can be distinguished by differences in their distribution profiles.

The fluorescence spectra were acquired at the same sites as the TA measurements. The ensemble average of 142 spectra, excluding one photobleached spectrum, is shown in **Figure 4a**. A narrow main peak and a broad tail were observed around 740 nm and above 770 nm, respectively. The spectral shape, especially the *R* value defined as the intensity ratio of the main peak to the tail, varied among individual aggregates at distinct sites (**Figure 4b**, red). The *R* reflects the exciton states in J-aggregates and is considered to be proportional to the exciton coherence length.^38-40^ The main peak and the broad tail of each spectrum were sufficiently reproduced with one and two Gaussian components, respectively (**Figure 4b**, blue, and see **Supporting Information** Section 9). The *R* was calculated by dividing the area of the main peak by that of the tail. Additionally, the fluorescence intensity *F* was calculated by integrating each fluorescence spectrum over the entire wavelength range. By dividing *F* by total Δ*A*, an effective fluorescence efficiency, denoted as *Φ*, was estimated. Thus, we obtained six photophysical parameters, including *τ*_1_, *τ*_2_, total Δ*A, F, Φ*, and *R*, for each measurement site.

**Figure 4.**
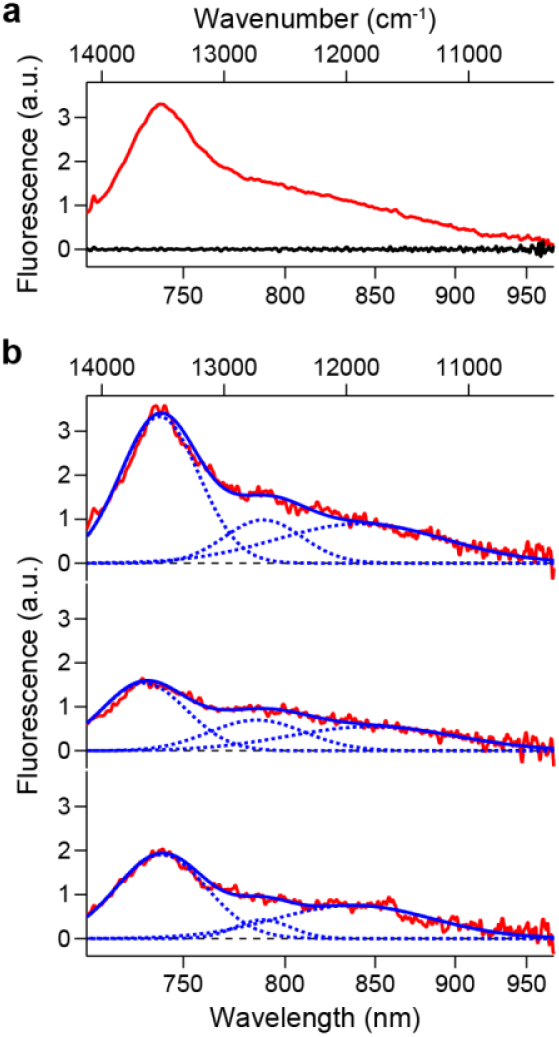
Fluorescence spectra of the Zn-HM aggregates. (a) Ensemble-averaged fluorescence spectrum. The black line indicates the background spectrum measured in a dark region of the fluorescence image. (b) Fluorescence spectra of individual aggregates at three distinct sites. The blue solid line represents the fitted curve, which is composed of three Gaussian components shown as blue dotted lines.

To photophysically characterize the C_B_ and C_N_ components in the *τ* distribution (**Figure 3c**), we examined 2D distributions constructed by plotting total Δ*A, F, Φ*, or *R* against *τ*_1_ and *τ*_2_ (**Figure 5**). The 2D histograms of total Δ*A* versus *τ*_1_ and *τ*_2_ were fitted with the sum of two 2D Gaussian functions representing the C_B_ and C_N_ components (**Figure 5a**). From this fit, the center and width of the total Δ*A* distribution and the correlation coefficient *r* between total Δ*A* and *τ* were estimated for the C_B_ and C_N_ components, respectively, as shown in **Figure 5b** (see **Supporting Information** Section 10). The total Δ*A* of C_N_ was 2.7 times that of C_B_, with a statistically significant difference (*p* < 10^−32^) (**Figure 5b**, red). C_B_ showed no significant correlation between *τ* and total Δ*A* (*r* = 0.08, *p* ≈ 0.3), whereas C_N_ showed a significant positive correlation (*r* = 0.42, *p* < 10^−4^) (**Figure 5b**, blue).

**Figure 5.**
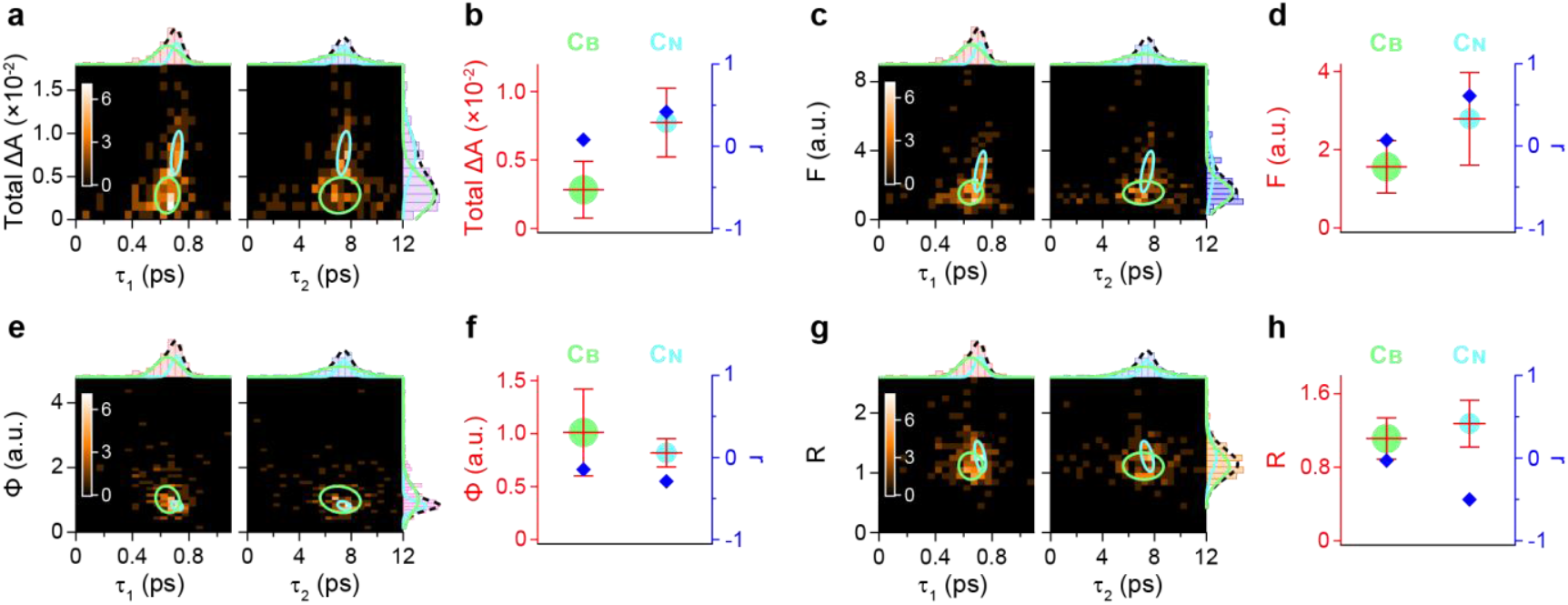
(a, c, e, g) 2D histograms of total Δ*A* (a), *F* (c), *Φ* (e), and *R* (g) versus *τ*_1_ (left panel) and *τ*_2_ (right panel). Circles indicate the contour at 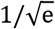 of the peak amplitude of the C_B_ (green) and C_N_ (cyan) components, identified by fitting with two-component 2D Gaussian functions. Projections of the distributions of each component onto the *τ*_1_, *τ*_2_, and corresponding parameter axes are shown in the same colors at the top of the left and right panels and at the right of the right panel, respectively. The dashed black lines indicate the summed distributions along each axis. (b, d, f, h) The mean values of total Δ*A* (b), *F* (d), *Φ* (f), and *R* (h) are indicated by red bars. The 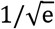 full widths are represented by the lengths of the vertical bars. The correlation coefficient *r* between *τ* and the corresponding parameter is indicated by blue diamonds. The areas of the green and cyan circles indicate the populations of the C_B_ and C_N_ components, respectively.

Applying the same 2D distribution analysis to *F, Φ*, and *R* (**Figures 5c–h**) revealed distinct trends between the C_B_ and C_N_ components. For C_N_, F was 1.8 times that of C_B_ (*p* < 10^−13^) and was positively correlated with *τ* (*r* = 0.61, *p* < 10^−9^), whereas F for C_B_ showed almost no correlation (*r* = 0.07, *p* ≈ 0.4) (**Figure 5d**). By contrast, *Φ* for C_N_ was 0.8 times that of C_B_ (*p* < 10^−7^) and showed a negative correlation with *τ* (*r* = −0.29, *p* < 10^−2^), while *Φ* for C_B_ exhibited a weaker correlation with marginal significance (*r* = −0.15, *p* ≈ 0.05) (**Figure 5f**). Likewise, *R* showed a statistically significant (*p* < 10^−5^) but small (1.1-fold) difference between C_B_ and C_N_, and *R* for C_N_ again showed a negative correlation with *τ* (*r* = −0.50, *p* < 10^−6^), whereas *R* for C_B_ showed no significant correlation (*r* = –0.03, *p* ≈ 0.7) (**Figure 5h**).

Decay time constants, *τ*_1_ and *τ*_2_, were estimated from the TA signals measured at distinct sites within individual Zn-HM aggregates. The faster component *τ*_1_ of ∼0.7 ps was comparable to the sub-ps excitation relaxation previously observed in TA measurements of porphyrin J-aggregates.^41^ In contrast, *τ*_2_ was ∼7.5 ps. Considering that time-resolved fluorescence measurements showed a decay faster than the ∼40-ps IRF (see **Supporting Information** Section 11), *τ*_2_ likely reflects depopulation of the excited state to the ground state or a non-emissive intermediate state. While a long-lived (ns) charge-transfer intermediate has been reported,^42^ nearly all of the TA signal amplitude decayed within 12 ps (**Figure 2h**), indicating recovery of ground-state absorption on the ps timescale and thereby excluding any substantial contribution from such long-lived intermediates. Therefore, *τ*_2_ was attributed to ground-state recovery. While ground-state recovery can proceed via radiative or non-radiative pathways, the lifetime of ∼7.5 ps is too short to be attributed to radiative decay. Although superradiance can accelerate emission from J-aggregates,^40^ the coherence length at room temperature is limited to a few to several tens of molecules,^38,43-44^ which is unlikely to account for the observed ∼7.5-ps lifetime. Accordingly, we assigned *τ*_2_ to non-radiative relaxation from the excited state to the ground state.

The 2D distribution analyses enabled us to characterize the C_B_ and C_N_ components by estimating total Δ*A*, fluorescence intensity *F*, fluorescence efficiency *Φ*, and fluorescence peak intensity ratio *R*, as well as their correlation coefficients *r* with *τ* (**Figure 5**). These observables reflect excitation dynamics in J-aggregates, particularly the properties of coherent excitonic domains in which excited states are delocalized over multiple molecules.^40^ Accordingly, our results can be explained by a heterogeneous aggregate model that incorporates exciton domains within J-aggregates, as described below (see **Figure S12** and **Supporting Information** Section 12).

C_N_ exhibited narrower *τ*_1_ and *τ*_2_ distribution widths than C_B_, which were 0.48 and 0.32 times those of C_B_, respectively (**Figure 3c** and **Table S3**). These results suggest that the C_N_ component arises from highly ordered J-aggregates formed through relatively homogeneous intermolecular interactions, because the time constants for non-radiative relaxation in J-aggregates depend on the strength of intermolecular coupling.^45^ C_N_ also showed a 2.7-fold larger total Δ*A* than C_B_ (**Figure 5b**, red), likely indicating a higher molecular packing density.

Even within the highly ordered C_N_ component, *τ* and other parameters exhibited a finite distribution width, suggesting that the aggregate structure was not fully uniform. A distinguishing feature of C_N_ was that the *τ* distribution was correlated with the distributions of other parameters. Specifically, *τ* and total Δ*A* were positively correlated (*r* = 0.42) (**Figure 5b**, blue), such that larger total Δ*A* corresponded to longer non-radiative relaxation time constants. Given theoretical predictions that a longer excitonic coherence length prolongs the non-radiative relaxation time constant,^43^ we infer that at sites with large total Δ*A*, a higher molecular packing density likely promotes the formation of excitonic states with relatively long coherence lengths, thereby lengthening the non-radiative relaxation time constant (**Figure S12a**).

Total Δ*A* primarily reflects the exciton domains that absorb the incident light. Meanwhile, emissive exciton domains can be assessed using the fluorescence intensity *F* and the fluorescence efficiency *Φ*. C_N_ exhibited an *F* value that was 1.8-fold larger than that of C_B_ (**Figure 5d**, red), consistent with its larger total Δ*A* because fluorescence output scales with absorbed light. By contrast, *Φ* differed only slightly between C_N_ and C_B_ (**Figure 5f**, red).

As observed for *τ* and total Δ*A, τ* and *F* in C_N_ were also positively correlated (*r* = 0.61) (**Figure 5d**, blue). In contrast, *Φ*, defined as the ratio of *F* to total Δ*A*, showed a weak negative correlation with *τ* (*r* = −0.29) (**Figure 5f**, blue), suggesting that *Φ* decreases with increasing *τ*. This trend implies a negative correlation between *Φ* and the coherence length in light-absorbing exciton domains. However, in J-aggregates, increasing the excitonic coherence length is expected to prolong non-radiative relaxation and simultaneously accelerate radiative decay via superradiance, thereby enhancing *Φ*.^40,43,46^ Therefore, if exciton domains were identical on the absorption and emission sides, *τ* and *Φ* should be positively correlated, contrary to our observation.

To avoid this discrepancy, we propose that excitation energy diffuses among exciton domains and is eventually trapped at the lowest-energy domain, from which fluorescence is emitted (**Figure S12b**). In this model, *Φ* increases with the coherence length of the emissive exciton domain. Concomitantly, the peripheral exciton domains, which absorb light, become relatively smaller and their coherence lengths shorten, reducing their non-radiative lifetimes. Consequently, the observed *τ* decreases. This inverse relationship between *τ* of the absorptive domain and the coherence length of the emissive domain explains the negative correlation between *τ* and *Φ* (**Figure 5f**, blue). Moreover, at loosely packed sites with smaller total Δ*A*, steric constraints and thus structural distortions are likely reduced, allowing the emissive domain to grow larger. If this expansion is accompanied by a further reduction in the size of the peripheral domains, *τ* would decrease, which provides a consistent interpretation of the positive correlation between *τ* and total Δ*A* (**Figure 5b**, blue).

Regarding the emissive exciton domain, the fluorescence spectrum provides further insight. For J-aggregates whose fluorescence spectra can be decomposed into a short-wavelength main peak and a long-wavelength tail (**Figure 4b**), the intensity ratio *R* (main peak/tail) is expected to be proportional to the excitonic coherence length.^38-40^ Accordingly, based on our interpretation in the preceding paragraph that the coherence length of the emissive exciton domain is inversely related to *τ, τ* and *R* should be negatively correlated, consistent with the observed result for C_N_ (*r* = –0.50) (**Figure 5h**, blue).

The *τ* distributions in C_B_ were broader than those in C_N_ (**Figure 3c**), suggesting greater heterogeneity in intermolecular interactions than in C_N_. Additionally, C_B_ exhibited a smaller total Δ*A*, implying a lower molecular packing density (**Figure 5b**, red). Therefore, we interpret the C_B_ regions as sparsely populated areas in which local intermolecular distances and orientations are highly nonuniform, leading to substantial variability in intermolecular coupling. This variability corresponds to a wide spread in the coherence length of excitonic domains (**Figure S12d**).

In C_B_, while the *Φ* distribution was broader than that in C_N_, its mean value remained comparable (**Figure 5f**, red). Likewise, *R* showed a mean value similar to that of C_N_ (**Figure 5h**, red). These results indicate that the coherence lengths of the emissive exciton domains are comparable in C_B_ and C_N_, consistent with a model in which excitation energy migrates among domains and is trapped at the lowest-energy exciton domain with a large coherence length, from which fluorescence is emitted. In the regions where C_B_ was observed, pronounced local structural heterogeneity is inferred to be present even within the femtoliter-scale volume defined by the diffraction-limited focal spot of the microscope. Accordingly, each measurement can be regarded as a subensemble average over heterogeneous microdomains within the focal spot, which explains why no strong correlations were detected between *τ* and the other parameters in C_B_ (**Figures 5b, d, f, h**, blue).

In this work, we developed a highly sensitive TA microscope. We further established an analytical framework that disentangles excitation dynamics using not only the mean values of time constants but also their distribution profiles, enabling us to identify multiple kinetic components with nearly identical time constants that are difficult to distinguish in conventional ensemble measurements. We analyzed the Zn-HM aggregates that mimic BChl self-aggregates in chlorosomes and revealed how microscopic differences in aggregate structure contribute to excitation dynamics. In native chlorosomes, however, BChl aggregates are further organized into tubular and lamellar suprastructures, forming a light-harvesting system rich in structural disorder.^5-7^ Owing to their pronounced heterogeneity, the properties and functional roles of these suprastructures remain poorly understood. The heterogeneity-resolved TA spectroscopy reported here should provide a powerful approach to elucidate the design strategies of natural light-harvesting systems that have evolved, while embracing heterogeneity, to function without relying on protein scaffolds.

### Notes

The authors declare no competing financial interest.

## ACKNOWLEDGMENT

The authors thank Dr. Shin Ogasawara and Dr. Hitoshi Tamiaki at Ritsumeikan University for providing samples, and Dr. Ken-ichi Inoue, Dr. Yutaka Shibata, and Dr. Shen Ye at Tohoku University for sharing laboratory space and for helpful discussions. This work was supported by Grants-in-Aid for Scientific Research (Nos. 25H00983, 23K23298, and 19H02665 to T.K.), JST-PRESTO (JPMJPR18G7 to T.K.), JST-FOREST (JPMJFR223M to T.K.), and JST-SPRING (JPMJSP2104 to S.A.). T.K. was supported by Yoshinori Ohsumi Fund for Fundamental Research, Asahi Glass Foundation, Research Foundation for Opto-Science and Technology, Murata Science Foundation, and Leading Initiative for Excellent Young Researchers (LEADER) from MEXT. S.M. was supported by The Hibi Science Foundation and JGC-S Scholarship Foundation.

## Notes

### Competing Interest Statement

The authors have declared no competing interest.

