## Supporting Information for "Heterogeneity-Resolved Ultrafast Transient Absorption Spectroscopy of Single Supramolecular Light-Harvesting Antennas"

**Supporting Information Section 1: Degrees of polarization of pump and probe beams**

The output of the laser source was linearly polarized. Both the pump and probe beams were passed through quarter-wave plates after being separated by a beam splitter (**Figure 1**). To quantify the degree of polarization, the beam was passed through a linear polarizer, and the transmitted light intensity was measured while rotating the polarizer. From the maximum ($I_{\max}$) and minimum ($I_{\min}$) intensities, the degree of polarization, defined as $(I_{\max} - I_{\min})/(I_{\max} + I_{\min})$, was calculated to be 0.09 and 0.25 for the pump and probe beams, respectively. Reducing the degree of linear polarization (i.e., approaching circular polarization) decreases the influence of sample orientation.

**Supporting Information Section 2: Sensitivity of the developed transient absorption (TA) microscope**

The minimum detectable $\Delta A$, defined as the measurement sensitivity, was evaluated as follows. As shown in **Figure S1a**, two band-pass filters, BPF1 and BPF2, with distinct spectral ranges were installed in the pump and probe beam paths, respectively. An optical chopper placed in the pump beam path modulated its intensity. This modulated light was combined with the unmodulated light in the probe beam path into a common path by a beam splitter. Subsequently, the combined beam was separated into two distinct beams, the signal and reference beams, by the second beam splitter. The modulated light included in the reference beam was removed by an optical filter. Consequently, only the unmodulated light was detected by the reference channel of the balanced detector. Meanwhile, the signal beam was focused by the objective, reflected by a mirror mounted on a coverslip, and collected by the same objective, as illustrated in the inset of **Figure 1**. The beam then passed through the second beam splitter and was eventually detected by the signal channel of the balanced detector. As a result, both the modulated and unmodulated light were present in the output of the signal channel.

The intensity of the unmodulated light in the signal beam was defined as $P_0$, which was measured while balancing the signal and reference beam intensities. The intensity of the modulated light in the signal beam was defined as $P_M$, which was selectively measured by lock-in detection. Here, $P_0$ corresponds to the steady-state intensity of the signal beam after being transmitted through the sample, whereas $P_M$ represents the intensity modulated by the pump beam, i.e., a pseudo TA signal. Therefore, the absorbance change $\Delta A$ can be calculated as $\Delta A = -\log(1 + P_M/P_0)$. Importantly, the experimentally obtained $P_0$ and $P_M$ inevitably include noise arising from the measurement apparatus and the environment. For example, the signal beam contains noise due to instabilities in microscope components such as the sample stage, as well as shot noise from the balanced photodiode. The sensitivity limit of the instrument is determined by these noise levels.

To experimentally evaluate the measurement sensitivity, we attenuated $P_M$ using neutral-density (ND) filters to progressively reduce $|\Delta A|$ and recorded the output voltage of the lock-in amplifier at each $|\Delta A|$. By comparing the output voltage with the noise voltage level, we estimated the detection limit as the $|\Delta A|$ value at which the signal-to-noise ratio (SNR) was equal to 1. **Figure S1b** shows the relationship between $|\Delta A|$ and the lock-in amplifier output voltage under various conditions, in which $P_0$, the time constant of the lock-in amplifier ($\tau_L$), and the wavelength of the modulated light ($\lambda_M$) were varied. The optical filter sets used for each condition are summarized in **Figure S1c**. The output voltage was acquired continuously with an acquisition interval of 0.2 s, and the signal intensity was obtained by averaging over 1 min. Under condition (i) ($P_0$ = 81 μW, $\tau_L$ = 30 ms, and $\lambda_M$ = 719–741 nm), the signal intensity decreased linearly as $|\Delta A|$ was reduced by attenuating $P_M$ (**Figure S1b**, red). The red dotted line indicates the signal intensity corresponding to SNR = 1, for which the noise level used to calculate the SNR was defined as the standard deviation of data acquired over 1 min in the absence of the modulated light ($P_M = 0$). Therefore, the intersection between the experimental data and the red dotted line represents the detection limit for $|\Delta A|$ at SNR = 1, which is defined as the measurement sensitivity. Under condition (i), the sensitivity was estimated to be ~$2\times10^{-7}$ (**Figure S1b**, red).

The absorbance of a single molecule of chlorophyll (Chl) *a* is given by $(\sigma/S)\cdot(1/\ln 10)$, where $\sigma$ is the absorption cross-section of Chl *a* and $S$ is the illuminated area. The molar extinction coefficient of Chl *a* is $8.13\times10^4$ $M^{-1}$ $cm^{-1}$,[1] from which $\sigma$ was calculated to be $3.11\times10^{-20}$ $m^2$. Using the full width at half maximum (FWHM) of the absorption focal spot along the *X*- and *Y*-axes as the diameter of the illuminated area, $S$ was calculated to be $6.41\times10^{-14}$ $m^2$. These values of $\sigma$ and $S$ yielded an absorbance of $2.11\times10^{-7}$ for a single Chl *a* molecule. This value is comparable to the sensitivity under condition (i), indicating that the developed TA microscope approaches the sensitivity required to detect absorbance changes at the single-molecule level.

Under condition (ii) in which $P_0 = 4$ μW, $\tau_L = 100$ μs, and $\lambda_M = 765$–775 nm, the sensitivity was on the order of $10^{-6}$ (**Figure S1b**, blue). Additionally, under condition (iii) in which $P_0 = 2$ μW, $\tau_L = 10$ μs, and $\lambda_M = 746$–755 nm, we estimated the sensitivity to be on the order of $10^{-4}$ (**Figure S1b**, green). In these measurements, the lock-in amplifier output was acquired continuously with an acquisition interval of 0.13 s. As described above, the sensitivity depends on $P_0$, $\tau_L$, and $\lambda_M$. Therefore, the measurement conditions should be optimized for each specific experiment.

The present TA measurements were performed under conditions nearly identical to condition (iii), i.e., $P_0 = 2$ μW (corresponding to ~14 μW just before the objective), $\tau_L = 10$ μs, and $\lambda_M = 746$–755 nm. The measured output voltage was in the range below $10^{-3}$ V, where the signal intensity and $|\Delta A|$ exhibited a linear relationship (**Figure S1b**, green). Therefore, using the parameters obtained from the linear fit, the output voltage of the lock-in amplifier was converted to $\Delta A$. The amplitudes of all TA signals in **Figures 2f, 2h, S3**, and **S4** were displayed as $\Delta A$.

a

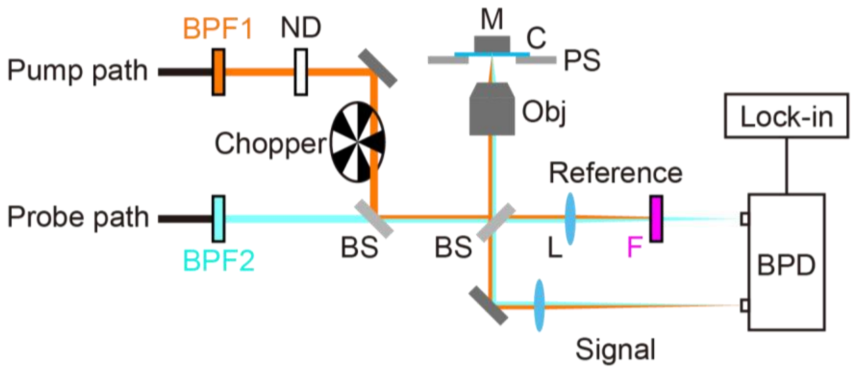


b

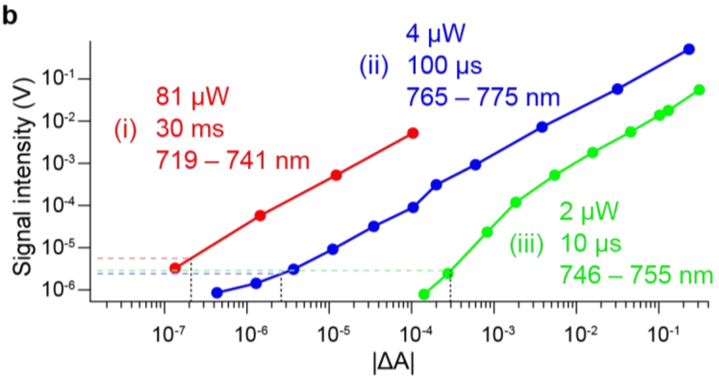


c

| | BPF1 | BPF2 | F |
|---|---|---|---|
| (i) | 719 – 741 nm (FF01-720/24-25)[a] | 765 – 775 nm (FBH770-10)[b] | >746 nm (FELH0750)[b] |
| (ii) | 765 – 775 nm (FBH770-10)[b] | 719 – 741 nm (FF01-720/24-25)[a] | <748 nm (FESH0750)[b] |
| (iii) | 746 – 755 nm (FBH750-10)[b] | 802 – 880 nm (FF01-793/LP-25)[a] | 802 – 880 nm (FF01-793/LP-25)[a] |

[a, b]: model number of the filters of Semrock and Thorlabs, respectively

**Figure S1.** Sensitivity assessment of the developed TA microscope. (a) Schematic of the optical setup employed to assess the measurement sensitivity. BPF: band-pass filter; ND: neutral-density filter; BS: beam splitter; Obj:

objective; L: lens; F: optical filter; BPD: balanced photodiode; Lock-in: lock-in amplifier; M: mirror; C: coverslip; and PS: piezo stage. (b) Relationship between the magnitude of the absorbance change $|\Delta A|$ and the lock-in amplifier output voltage under different conditions (i)–(iii), shown in red, blue, and green, respectively. The values of $P_0$, $\tau_L$, and $\lambda_M$ under each condition are given in the panel. The output voltage and $|\Delta A|$, at which the SNR was equal to 1 under each condition, are indicated by horizontal and vertical dotted lines, respectively. (c) Optical filter sets used under each condition, corresponding to BPF1, BPF2, and F in (a).

**Supporting Information Section 3: Spatial resolution of fluorescence and absorption microscopy**

Fluorescence and absorption imaging of individual chlorosomes was performed to estimate the spatial resolution of the microscope. Chlorosomes are light-harvesting antennas of green sulfur bacteria and typically form vesicles with dimensions of 70–180 × 30–60 × 10–20 nm, containing bacteriochlorophyll (BChl) *c*, *d*, and *e*.[2]

To assess the spatial resolution along the *X*- and *Y*-axes, fluorescence and absorption images were acquired in the focal (*XY*) plane with an excitation fluence of 5–7×$10^{14}$ photons $\text{pulse}^{-1}$ $\text{cm}^{-2}$ at 719–741 nm (**Figures S2a, d**, respectively). In these measurements, the long-pass filters placed in front of the balanced detector (**Figure 1**) were removed. By fitting the fluorescence spot in the *XY* plane to a two-dimensional (2D) Gaussian function (**Figure S2a**), FWHMs along the *X*- and *Y*-axes were estimated to be 309 and 257 nm, respectively, corresponding to the spatial resolution for fluorescence measurements (**Figures S2b, c**). Similarly, by fitting the absorption spot in the *XY* plane to a 2D Gaussian function (**Figure S2d**), FWHMs defining the spatial resolution for absorption measurements were estimated to be 336 and 243 nm along the *X*- and *Y*-axes, respectively (**Figures S2e, f**).

Furthermore, we acquired fluorescence and absorption images along the optical (*Z*) axis (**Figures S2g, i**, respectively). As the distance between the objective and the sample was increased by moving the piezo stage along the *Z*-axis, the light reflected by the mirror placed on the sample coverslip was first focused onto the sample, as shown in the inset of **Figure 1**. Upon further increasing the distance, the light was temporarily defocused, but then refocused onto the sample directly without being reflected by the mirror. Consequently, two spots were observed along the *Z*-axis in both fluorescence and absorption images (**Figures S2g, i**, respectively). The separation distance between these two focal planes was approximately 2 μm, half of which (≈ 1 μm) corresponds to the distance between the sample coverslip and the mirror placed on it. In the measurements of the Zn-HM aggregates, the coverslip–mirror distance was estimated to be approximately 8 μm, and all measurements were performed at the lower *Z* focal plane.

The fluorescence intensity profile along the *Z*-axis was fitted with a one-dimensional (1D) Gaussian function (**Figure S2h**), yielding a FWHM of 1031 nm, corresponding to the spatial resolution along the optical axis for fluorescence measurements. For absorption measurements, the spatial resolution along the *Z*-axis was estimated to be 1145 nm (**Figure S2j**). These fluorescence and absorption profiles were fitted simultaneously, sharing a common central *Z* position as a global parameter.

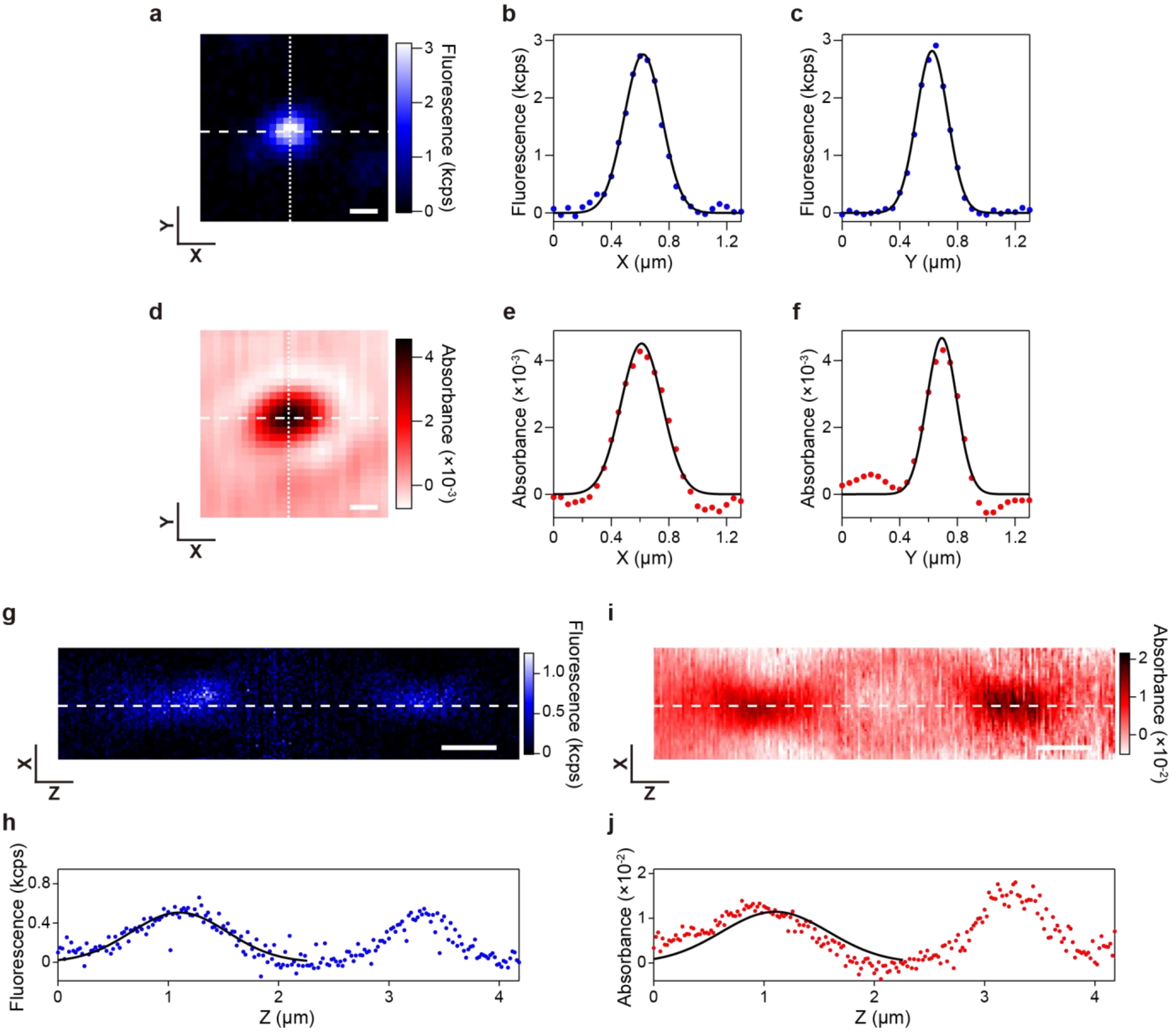


**Figure S2.** Fluorescence and absorption images along the *X*-, *Y*-, and *Z*-axes. (a, d) Fluorescence (a) and absorption (d) images of a single chlorosome in the focal (*XY*) plane. The scale bar represents 0.2 μm. (b, c, e, f) Line profiles of fluorescence intensity (b, c) and absorbance (e, f) along the *X*- and *Y*-axes, as represented by dashed and dotted lines, respectively, in (a) and (d). Solid lines indicate the fitted curves. (g, i) Fluorescence (g) and absorption (i) images in the optical plane (*XZ* plane). The scale bar represents 0.5 μm. (h, j) Line profiles of fluorescence intensity (h) and absorbance (j) along the *Z*-axis, as represented by a dashed line in (g) and (i). Solid lines indicate the fitted curves.

**Supporting Information Section 4: Sample preparation**

Zinc 3-hydroxymethyl pyropheophorbide-*a* (Zn-HM) was synthesized according to previous reports.[3-4] Zn-HM was dissolved in $CHCl_3$ (100 μL), and then hexane (1900 μL) was added to the $CHCl_3$ solution to give a dispersion of assembled Zn-HM in 1% $CHCl_3$-hexane (2 mL). The dispersion was allowed to stand for 1 week to prepare the sample for microscopy measurements.

The Zn-HM aggregates were fixed on a coverslip by drop casting. Subsequently, a 40 mM Tris-HCl (pH 8) buffer containing 1% polyvinyl alcohol and an oxygen-scavenging system of 1 U/mL protocatechuate 3,4-dioxygenase (Oriental Yeast) and 2.5 mM protocatechuic acid was spin-coated onto the sample at 3000 rpm for 1 min. Moreover, an Ag mirror (PF03-03-P01, Thorlabs), which was spin-coated with 1 µL of immersion oil at 6000 rpm for 1 min to form a thin oil layer on its surface, was placed on the sample. The coverslip–mirror distance was estimated to be approximately 8 µm.

**Supporting Information Section 5: Estimation of the number of decay components in TA signals and of the instrument response function (IRF)**

The TA signal was fitted with an exponential decay function convolved with a Gaussian function corresponding to the instrument response function (IRF), given by:

$$\mathrm{f}(t) = y_0 + \frac{\sqrt{\pi}\, t_\mathrm{w}}{2} \sum_{i=1}^{N} A_i \exp\left(\frac{t_0 - t}{\tau_i} + \frac{t_\mathrm{w}^2}{4\tau_i^2}\right) \mathrm{erfc}\left(\frac{t_0 - t + \frac{t_\mathrm{w}^2}{2\tau_i}}{t_\mathrm{w}}\right), \quad (1)$$

where $y_0$ is the baseline, $t_\mathrm{w}$ is the half width at 1/e of the maximum of the IRF, $N$ is the number of decay components, $t_0$ is the offset along the $t$-axis, and $A_i$ and $\tau_i$ are the amplitude and the time constant of decay component $i$, respectively. erfc denotes the complementary error function, expressed as follows:

$$\mathrm{erfc}(t) = \frac{2}{\sqrt{\pi}} \int_t^{\infty} \exp(-z^2)\, \mathrm{d}z\ . \quad (2)$$

To determine the number of decay components and the width of the IRF, we analyzed an ensemble-averaged TA signal obtained by averaging TA signals acquired at distinct sites (**Figure S3**, red). Specifically, among the 159 signals acquired in total, we averaged 145 signals whose amplitude, defined as the mean of five data points around the peak at $t = 0$ ps, exceeded three times the noise level, given by the standard deviation of the background signal measured at a non-fluorescent site (**Figure 2f**, black). This ensemble-averaged TA signal was fitted using eq. (1) with different $N$ values. First, a one-component fit ($N = 1$) did not reproduce the experimental result well (**Figure S3**, cyan). By contrast, a two-component fit ($N = 2$) provided a reasonable result (**Figure S3**, blue), reducing the $\chi^2$ value, defined as the sum of the squared residuals between the data and the fitted curve, to 8% of that obtained with the one-component fit (**Table S1b**). Even when the number of decay components was increased to 3 ($N = 3$), the fitted result was almost the same as that obtained with the two-component fit (**Figure S3**, green). The $\chi^2$ value decreased to 3% of that obtained with the one-component fit, indicating no substantial improvement compared with the two-component fit (**Table S1c**). Therefore, we concluded that two decay components are sufficient to interpret the TA signals. From the two-component fit, the FWHM of the IRF was estimated to be 177 fs (**Table S1b**).

Using eq. (1) with $N = 2$ and an IRF FWHM of 177 fs, TA signals from individual Zn-HM aggregates acquired at distinct measurement sites were each fitted. The 143 signals (out of 159) for which the fitted curves showed a maximum amplitude exceeding three times the noise level, defined as the standard deviation of the background signal, were used for subsequent statistical analyses.

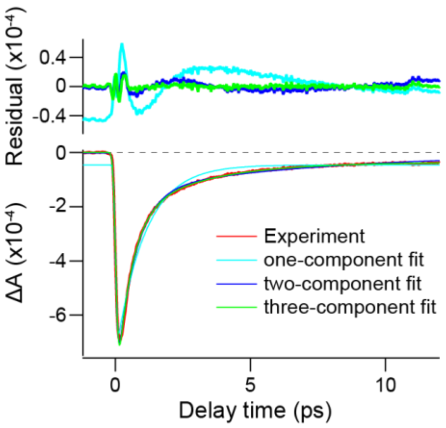


**Figure S3.** Ensemble-averaged TA signal fitted with multiple exponential components. The bottom panel shows the ensemble-averaged TA signal obtained by averaging 145 signals from individual Zn-HM aggregates measured at distinct sites (red), together with the fits obtained using one-, two-, and three-component exponential decay functions (cyan, blue, and green, respectively). The top panel shows the corresponding residuals. The ensemble-averaged TA signal (red) and the two-component fit (blue) are replotted in **Figure 2f** with the vertical axis rescaled.

**Table S1.** Parameters estimated from one-, two-, and three-component exponential fits to the ensemble-averaged TA signal.

| | | $\tau$ (ps) (relative $\Delta A$) | | | IRF (fs) | $\chi^2$ |
|---|---|---|---|---|---|---|
| | | component 1 | component 2 | component 3 | | |
| (a) | $N = 1$ | 1.1 (1.00) | - | - | 134 | $1.94\times10^{-7}$ (100%) |
| (b) | $N = 2$ | 0.7 (0.83) | 7.2 (0.17) | - | 177 | $0.15\times10^{-7}$ (8%) |
| (c) | $N = 3$ | 0.5 (0.72) | 2.7 (0.24) | 208.5 (0.04) | 187 | $0.06\times10^{-7}$ (3%) |

**Supporting Information Section 6: Temporal fluctuations in the TA signals**

**Figure S4** shows the time-series data of the TA signals at sites 1–3 in **Figure 2e**, which were acquired continuously at 0.25-s intervals and averaged over 2-s intervals. The TA signals at each measurement time were fitted with a two-component exponential function given by eq. (1) to quantify the temporal variations in $\Delta A$ and $\tau$ (**Figure S5**). At site 1, the mean and standard deviation of $\tau_1$ were 0.8 ps and 0.1 ps, respectively. The coefficient of variation, defined as the standard deviation divided by the mean, was 13% (**Figure S5a**, top panel, red). For $\tau_2$, the mean, standard deviation, and coefficient of variation were 8.8 ps, 2.2 ps, and 26%, respectively (**Figure S5a**, bottom panel, red). The coefficients of variation for both $\tau_1$ and $\tau_2$ were greater than the fitting inaccuracy for the estimated time constants, which was determined, as described in **Supporting Information Section 7**, to be less than 6% based on an SNR of around 10 obtained from each TA signal in the time series. Therefore, these temporal variations likely reflect fluctuations in excitation dynamics arising from dynamic disorder within the local measurement region. Distinct behaviors were observed at sites 2 and 3 (**Figures S5b, c**). Across the three sites, the mean time constants ranged from 0.5–0.8 ps for $\tau_1$ and 6.1–8.8 ps for $\tau_2$. These variations among sites suggest heterogeneity in excitation dynamics arising from static disorder. At site 2, photobleaching was observed as a progressive decrease in $\Delta A_1$ over time (**Figure S5b**, top panel, blue). In contrast, at sites 1 and 3, only a slight decrease in $\Delta A_1$ was observed, indicating that photobleaching was relatively slow (**Figures S5a, c**, top panel, blue). Thus, the timescale of photobleaching was also heterogeneous across sites.

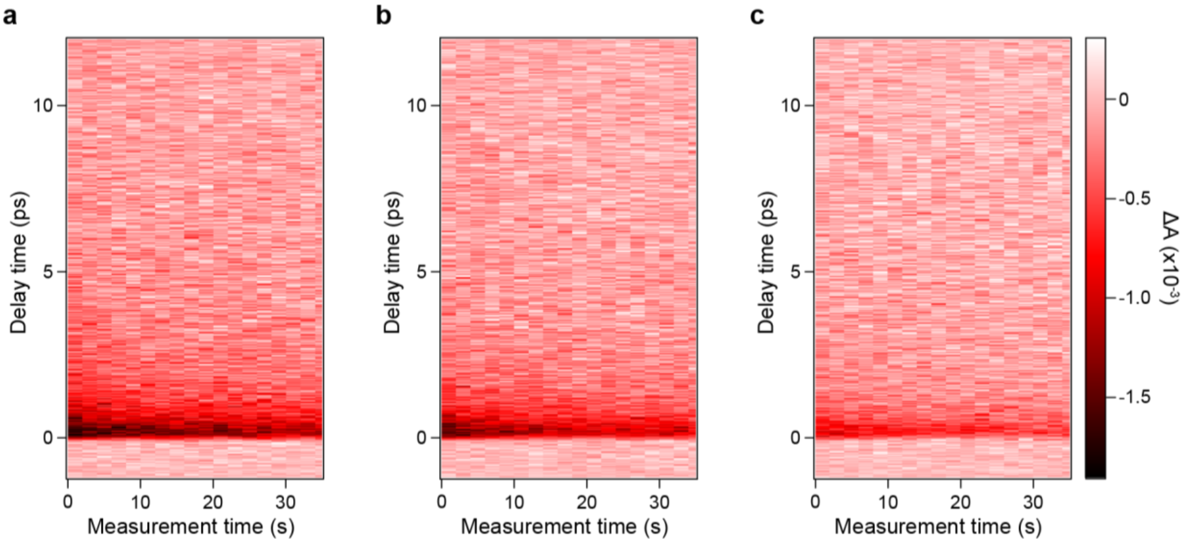


**Figure S4.** Time-series TA signals sequentially acquired from individual Zn-HM aggregates. TA signals were averaged over 2-s intervals. The resulting time series at sites 1, 2, and 3 in **Figure 2e** are shown in (a), (b), and (c), respectively.

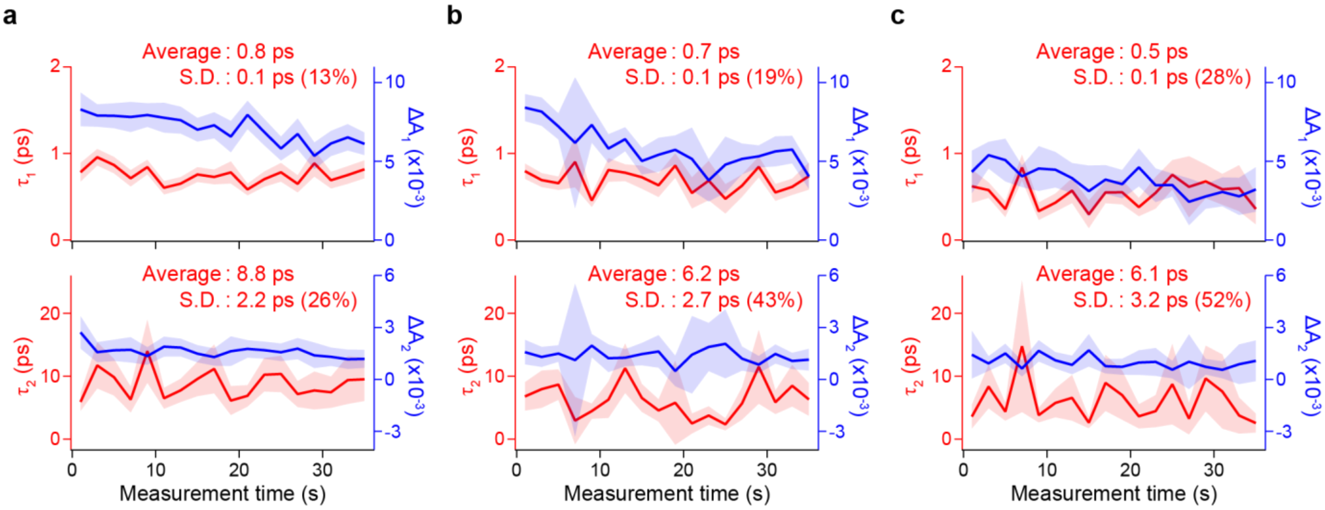


**Figure S5.** Temporal variations in TA kinetics of individual Zn-HM aggregates. Time-dependent changes in the time constants (red) and the magnitudes of the absorbance changes (blue) at sites 1, 2, and 3 in **Figure 2e** are shown in (a), (b), and (c), respectively. These values were estimated by fitting each TA signal in the time series displayed in **Figure S4** with a two-component exponential function. The top panel corresponds to $\tau_1$ and $\Delta A_1$ of the fast decay component, and the bottom panel corresponds to $\tau_2$ and $\Delta A_2$ of the slow decay component. The means, standard deviations (S.D.), and coefficients of variation, defined as S.D./mean, for $\tau_1$ and $\tau_2$ are indicated in each panel. The shaded areas represent ± one standard error for the fit at each time point. Traces in (a) are replotted in **Figure 2g** with the vertical axis rescaled.

**Supporting Information Section 7: Reliability of time-constant estimates exceeding the measurement time window**

Among the fitted signals, 14% (20 out of 143) yielded time constants that exceeded the 12-ps time window of the TA measurements. The reliability of these long time constants was assessed by a simulation analysis, as described below. First, we generated synthetic exponential decay curves assuming a single-exponential decay over the 12-ps window, with the decay time constant $\tau$ set in the range of 5–40 ps. The FWHM of the IRF was fixed at 177 fs, as determined experimentally (**Table S1b**). The decay curves were simulated with the SNR set to 23, corresponding to the median SNR of the experimental data used for our analysis (**Figure S6a**, orange), and were then fitted with a single-exponential function to estimate the time constant (**Figure S6a**, blue). From 1,000 trials of the simulation and fits, we obtained the standard deviation of the estimated time constants. The coefficient of variation, calculated by dividing the standard deviation by the set $\tau$, was defined as the relative fitting error and plotted as a function of the set $\tau$ in the range of 5–40 ps (**Figure S6c**, orange). The relative fitting error remained nearly constant for $\tau < 12$ ps, but increased as the set $\tau$ increased beyond 12 ps. When the SNR was set to 3, corresponding to the lower limit of the experimental data used for our analysis, the fitting accuracy appeared to decline (**Figure S6b**). Meanwhile, as in the case of SNR = 23, the relative fitting error also increased for $\tau > 12$ ps (**Figure S6c**, pink). These results indicate that the reliability of time-constant estimates decreases for time constants exceeding the measurement time window. Accordingly, the statistical distribution analyses associated with the time constants, as shown in **Figures 3 and 5**, were performed using time constants of $\tau < 12$ ps.

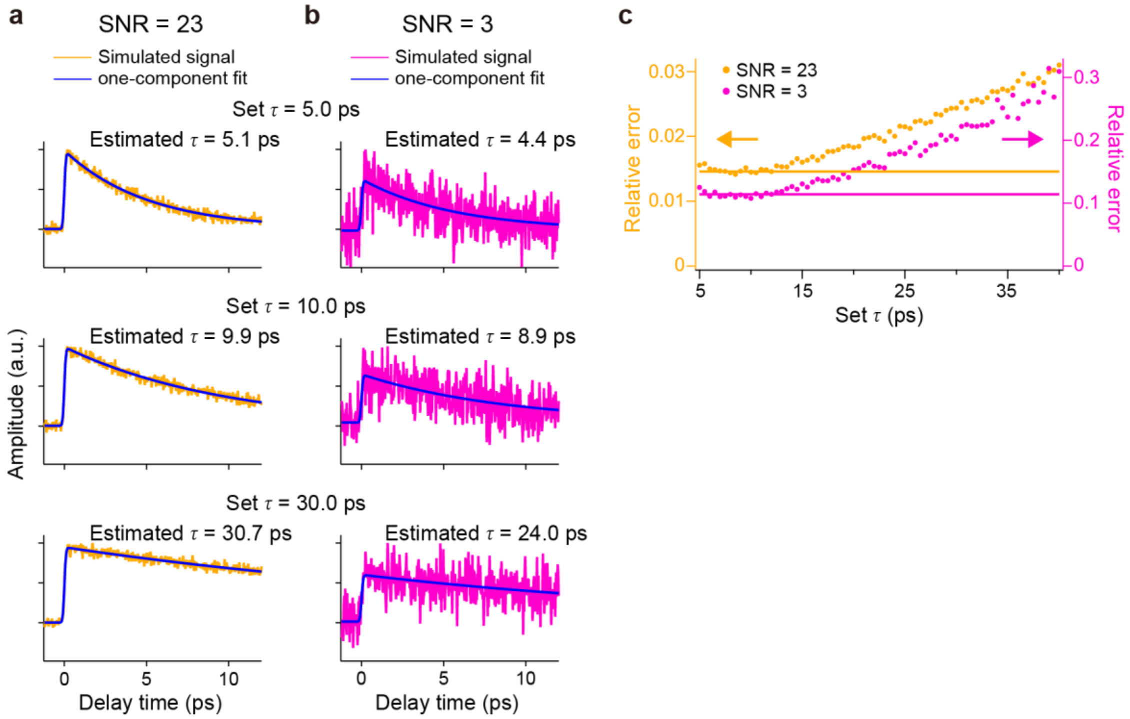


**Figure S6.** Reliability assessment of time-constant estimates. (a, b) Synthetic single-exponential decay signals generated over a 12-ps time window for SNRs of 23 (a) and 3 (b). Three representative decay signals calculated with $\tau$ set to 5, 10, and 30 ps are shown from top to bottom, respectively. Fitted curves are indicated by blue lines. The set and estimated $\tau$ values are annotated in each panel. (c) Relative fitting error estimated for SNRs of 23 (orange) and 3 (pink), plotted as a function of the set $\tau$. The error levels at a set $\tau$ of 12 ps, corresponding to the 12-ps time window, are indicated by horizontal lines for each SNR condition.

**Supporting Information Section 8: Fitting analysis of distributions of $\tau$ and other parameters**

**(i) Determination of the number of components in $\tau$ distributions**

**Figures 3a and 3b** show the distributions of $\tau_1$ and $\tau_2$, constructed from the fitted $\tau$ values obtained from 143 and 123 TA signals, respectively. Of the 143 signals, 20 exhibited $\tau_2$ values longer than the 12-ps measurement time window, and thus statistical analyses of $\tau_2$ were performed using the remaining 123 signals (see **Supporting Information Section 7**). The profiles of the $\tau_1$ and $\tau_2$ histograms clearly indicate contributions of multiple distribution components. Therefore, we performed a fitting analysis with a multi-component Gaussian function, given by:

$$\mathrm{f}(\tau) = \sum_{i=1}^{N} A_i \exp\left(-\left(\frac{\tau - \tau_{\mathrm{c}i}}{\tau_{\mathrm{w}i}}\right)^2\right), \tag{3}$$

where $N$ is the number of distribution components, and $A_i$, $\tau_{\mathrm{c}i}$, and $\tau_{\mathrm{w}i}$ are the amplitude, central position, and 1/e width for component $i$, respectively. The fitting results obtained using eq. (3) with different $N$ values from 1 to 3 are summarized in **Table S2**. As described below, the optimal $N$ for the $\tau_1$ and $\tau_2$ histograms was determined based on the $\chi^2$ values.

Fitting the histogram of $\tau_1$ with a one-component Gaussian function ($N = 1$) could not reproduce the distribution in the 0.3–0.5 ps range (**Figure S7a**, cyan and **Table S2a**). A two-component fit ($N = 2$) provided two peaks centered at 0.62 ps with a FWHM of 0.33 ps and at 0.69 ps with a FWHM of 0.12 ps, sufficiently reproducing the overall distribution profile including the 0.3–0.5 ps range (**Figure S7a**, blue and **Table S2b**). The $\chi^2$ value decreased to 19% of that obtained with the one-component fit. Increasing $N$ to 3 resulted in an additional sharp peak at 0.41 ps with a FWHM of 0.05 ps (**Figure S7a**, green and **Table S2c**). The FWHM of this third peak was unnaturally small, comparable to the bin size of 0.05 ps for the $\tau_1$ histogram, suggesting overfitting due to an excessively large $N$. Furthermore, the $\chi^2$ value, while decreasing to 11% compared to the one-component fit, showed no substantial improvement over the two-component fit. Accordingly, we concluded that $N = 2$ is optimal for the $\tau_1$ histogram.

The same analysis was performed on the $\tau_2$ histogram. As in the case of $\tau_1$, while a one-component fit ($N = 1$) failed to reproduce the tail of the peak (**Figure S7b**, cyan and **Table S2d**), a two-component fit ($N = 2$) yielded a reasonable profile composed of two peaks centered at 7.30 ps with a FWHM of 4.97 ps and at 7.42 ps with a FWHM of 1.39 ps (**Figure S7b**, blue and **Table S2e**). The $\chi^2$ value decreased to 56% relative to the one-component fit. By contrast, a three-component fit ($N = 3$) exhibited a signature of overfitting as a sharp peak at 5.57 ps with an unnaturally narrow FWHM of 0.52 ps, comparable to the bin size of 0.5 ps for the $\tau_2$ histogram (**Figure S7b**, green and **Table S2f**). The $\chi^2$ value decreased to 47% of that obtained with the one-component fit, which is comparable to the reduction achieved by the two-component fit (**Table S2f**). These results indicated that $N = 2$ is also optimal for the $\tau_2$ histogram, as in the case of $\tau_1$.

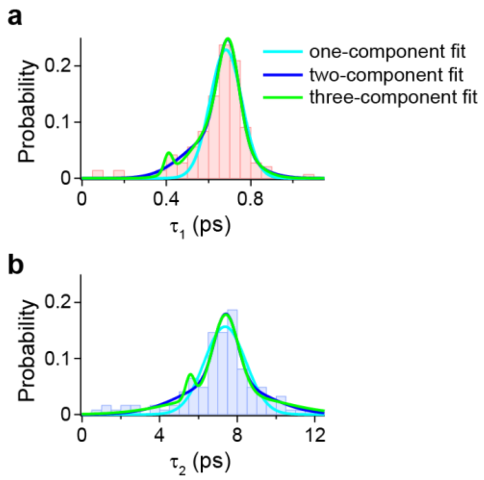


**Figure S7.** $\tau$ distributions fitted with Gaussian functions. Histograms of $\tau_1$ (a) and $\tau_2$ (b) were fitted with one-, two-, and three-component Gaussian functions (cyan, blue, and green, respectively). The bin sizes were 0.05 ps for $\tau_1$ and 0.5 ps for $\tau_2$. These histograms are replotted in **Figures 3a and 3b**, respectively.

**Table S2.** Parameters estimated from one-, two-, and three-component Gaussian fits to the $\tau_1$ and $\tau_2$ distributions.

| $\tau_1$ distribution | | | | | |
|---|---|---|---|---|---|
| | Number of components | Population (%) | $\tau$ (ps) | FWHM (ps) | $\chi^2$ |
| (a) | 1 | 100 | 0.68 | 0.18 | $5.2 \times 10^{-3}$ (100%) |
| (b) | 2 | 51<br>49 | 0.62<br>0.69 | 0.33<br>0.12 | $1.0 \times 10^{-3}$ (19%) |
| (c) | 3 | 3<br>57<br>40 | 0.41<br>0.65<br>0.70 | 0.05<br>0.27<br>0.11 | $0.6 \times 10^{-3}$ (11%) |

| $\tau_2$ distribution | | | | | |
|---|---|---|---|---|---|
| | Number of components | Population (%) | $\tau$ (ps) | FWHM (ps) | $\chi^2$ |
| (d) | 1 | 100 | 7.37 | 2.51 | $8.1 \times 10^{-3}$ (100%) |
| (e) | 2 | 63<br>37 | 7.30<br>7.42 | 4.97<br>1.39 | $4.5 \times 10^{-3}$ (56%) |
| (f) | 3 | 5<br>47<br>48 | 5.57<br>7.40<br>7.57 | 0.52<br>1.56<br>6.25 | $3.9 \times 10^{-3}$ (47%) |

**(ii) 2D distribution analysis**

The 2D histogram of $\tau_1$ versus $\tau_2$ shows a distribution along the diagonal (**Figure 3c**), reflecting a strong positive correlation and indicating that the $\tau_1$ and $\tau_2$ distributions are tightly coupled. As described above, because both the $\tau_1$ and $\tau_2$ histograms contain two distribution components (**Figure S7**), their 2D distribution can likewise comprise two components. Thus, we performed a fitting analysis with a function consisting of two 2D Gaussian components, given by:

$$f(\tau_1, \tau_2) = \sum_i^2 A_i \exp\left( -\frac{1}{2\,(1 - r_i^2)} \left( \left( \frac{\tau_1 - \tau_{c1,i}}{\tau_{w1,i}} \right)^2 + \left( \frac{\tau_2 - \tau_{c2,i}}{\tau_{w2,i}} \right)^2 - \frac{2 \cdot r_i \cdot (\tau_1 - \tau_{c1,i}) \cdot (\tau_2 - \tau_{c2,i})}{\tau_{w1,i} \cdot \tau_{w2,i}} \right) \right), \quad (4)$$

where $i$ indexes the two components and $A_i$ is the amplitude for component $i$. $\tau_{c1,i}$ and $\tau_{w1,i}$ ($\tau_{c2,i}$ and $\tau_{w2,i}$) represent the central position and $1/\sqrt{e}$ width for component $i$, respectively, along the $\tau_1$-axis ($\tau_2$-axis). $r_i$ is the correlation coefficient between $\tau_1$ and $\tau_2$ for component $i$. Fitting the 2D histogram with eq. (4) identified two distribution components with distinct widths, i.e., a broadly distributed component ($C_B$) and a narrowly distributed component ($C_N$) (**Figure 3c**, green and cyan, respectively). The population of each component was estimated by calculating the ratio of the distribution volumes of the two components, given by $2\pi \cdot A_i \cdot \tau_{w1,i} \cdot \tau_{w2,i} \cdot \sqrt{1 - r_i^2}$. The fitting results are summarized in **Table S3**.

**Table S3.** Parameters estimated from a two-component 2D Gaussian fit to the 2D distribution of $\tau_1$ versus $\tau_2$.

| | Population (%) | $\tau_1$ (ps) | FWHM (ps) | $\tau_2$ (ps) | FWHM (ps) | $r$ |
|---|---|---|---|---|---|---|
| $C_B$ | 67 | 0.6 | 0.20 | 7.1 | 3.65 | 0.86 |
| $C_N$ | 33 | 0.7 | 0.09 | 7.4 | 1.17 | 0.83 |

**(iii) Distributions of the expected time constant $\langle\tau\rangle$, total $\Delta A$, fluorescence intensity $F$, fluorescence efficiency $\Phi$, and fluorescence peak intensity ratio $R$.**

At each measurement site, the TA signal was fitted using eq. (1) to estimate two time constants ($\tau_1$ and $\tau_2$), two amplitudes ($\Delta A_1$ and $\Delta A_2$), and their total amplitude (total $\Delta A$). **Figure S8a** shows the histogram of the expected time constant $\langle\tau\rangle$, defined as $(\Delta A_1 \cdot \tau_1 + \Delta A_2 \cdot \tau_2)$ / total $\Delta A$, which was constructed with a bin size of 0.2 ps. The mean and median values are indicated by a black dotted line and a red solid line, respectively. Conventional ensemble measurements typically yield a value close to the mean of this distribution. In addition, the histogram of total $\Delta A$ was constructed with a bin size of $1.0\times10^{-3}$ (**Figure S8b**).

**Figure S8c** shows the histogram constructed with a bin size of 0.3 for the fluorescence intensity $F$, defined as the integrated intensity of the fluorescence spectrum acquired at each site (see **Supporting Information Section 9**). Whereas $F$ reflects the amount of light energy emitted as fluorescence, total $\Delta A$ corresponds to the amount of absorbed light energy. Therefore, the ratio ($F$ / total $\Delta A$) provides an effective fluorescence efficiency $\Phi$ at each site. The histogram of $\Phi$ constructed with a bin size of 0.1 is shown in **Figure S8d**, where $\Phi$ is normalized by its median value. Furthermore, we constructed the histogram of the fluorescence peak intensity ratio $R$ with a bin size of 0.1 (**Figure S8e**). $R$ is calculated as the ratio of the main peak area to the tail peak area in the fluorescence spectrum (see **Supporting Information Section 9**).

Each histogram spans a broad range and deviates from a simple Gaussian profile, indicating contributions from multiple distribution components. Using the 2D distribution analysis described in **Supporting Information Section 10**, the components can be distinguished and identified while linking all parameters to one another.

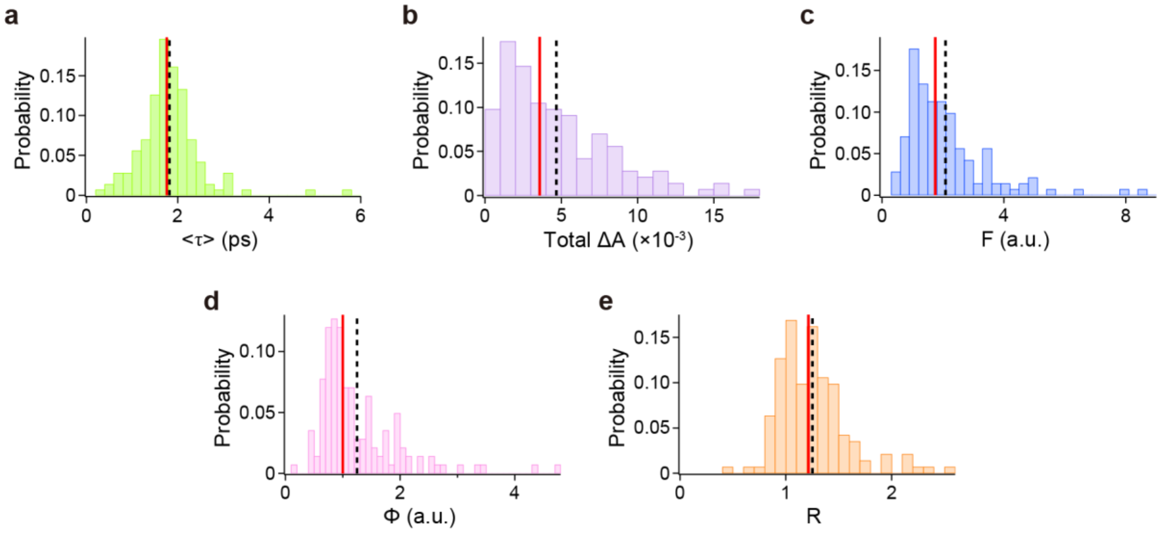


**Figure S8.** Distributions of multiple parameters. Histograms of the expected time constant $\langle\tau\rangle$ (a), total $\Delta A$ (b), fluorescence intensity $F$ (c), fluorescence efficiency $\Phi$ (d), and fluorescence peak intensity ratio $R$ (e) were constructed with bin sizes of 0.2 ps, $1.0\times10^{-3}$, 0.3, 0.1, and 0.1, respectively. The black dotted and red solid lines indicate the mean and median values of each distribution, respectively.

**Supporting Information Section 9: Fluorescence spectral analysis**

Fluorescence spectra of individual Zn-HM aggregates were acquired with an excitation wavelength of ~695 nm (**Figure 2d**, blue) and an excitation fluence of $4.0\times10^{13}$ photons pulse$^{-1}$ cm$^{-2}$, using an integration time of 60 s. The fluorescence wavelength was calibrated using the measured spectrum of a Ne lamp. The wavelength-dependent detection sensitivity was also calibrated by fitting the measured spectrum of a halogen lamp to Planck's law of black-body radiation.

The fluorescence spectra were acquired at the same sites as the TA measurements. Of the 143 spectra, 142 were averaged, excluding one photobleached spectrum. The ensemble-averaged spectrum exhibited a narrow main peak around 740 nm and a broad tail above 770 nm (**Figure S9**, red). The main peak was reproduced well by a Gaussian function centered at 13,509 cm$^{-1}$ with a FWHM of 803 cm$^{-1}$ (**Figure S9**, black dashed lines). In contrast, the spectral tail could not be reproduced by a single Gaussian function (**Figure S9a** and **Table S4a**). A two-component fit adequately reproduced the tail (**Figure S9b**), reducing the $\chi^2$ value to 27% of that obtained with the one-component fit (**Table S4b**). A three-component fit yielded no substantial improvement relative to the two-component fit (**Figure S9c**). The $\chi^2$ value decreased to 17% of that obtained with the one-component fit, comparable to that obtained with the two-component fit (**Table S4c**). Based on these results, we concluded that two Gaussian components were necessary to reproduce the tail. The tail intensity was defined as the summed area of the two Gaussian components, whereas the main peak intensity was defined as the area of the single Gaussian used to fit the main peak. The fluorescence peak intensity ratio $R$ was calculated by dividing the main peak intensity by the tail intensity. Moreover, the fluorescence intensity $F$ was calculated by integrating the spectral intensity over the entire wavelength range.

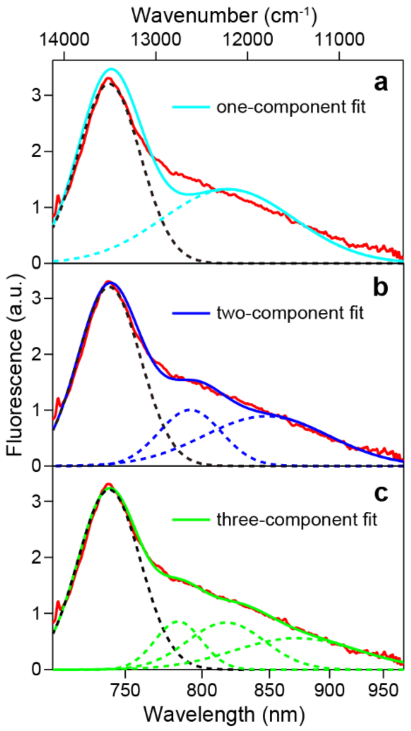


**Figure S9.** Ensemble-averaged fluorescence spectrum fitted with multiple Gaussian components. The ensemble-averaged fluorescence spectrum was obtained by averaging 142 spectra of individual Zn-HM aggregates measured at distinct sites (red). The spectral tail was fitted with one- (a), two- (b), and three-component (c) Gaussian functions, as indicated by dotted lines in cyan, blue, and green, respectively. The narrow main peak was fitted with the same single Gaussian component in all panels, indicated by a black dotted line. Solid lines show the sum of all components. The ensemble-averaged fluorescence spectrum (red) is replotted in **Figure 4a** with the vertical axis rescaled.

**Table S4.** Parameters estimated from one-, two-, and three-component Gaussian fits to the spectral tail of the ensemble-averaged fluorescence spectrum.

| | | Tail peak | | | |
|---|---|---|---|---|---|
| | Number of components | Population (%) | Central position ($cm^{-1}$) | FWHM ($cm^{-1}$) | $\chi^2$ |
| (a) | 1 | 46 | 12,211 | 1,686 | 58 (100%) |
| (b) | 2 | 17<br>30 | 12,628<br>11,799 | 798<br>1,604 | 16 (27%) |
| (c) | 3 | 11<br>17<br>19 | 12,758<br>12,238<br>11,471 | 601<br>991<br>1,602 | 10 (17%) |

### Supporting Information Section 10: 2D distribution analysis of $\tau$ versus other photophysical parameters

#### (i) Fitting procedure for 2D distribution analysis

In this study, we identified the $C_B$ and $C_N$ components in the $\tau_1$ and $\tau_2$ distributions (**Figure 3**). We also obtained distributions of four parameters, including the total absorbance change (total $\Delta A$), fluorescence intensity ($F$), fluorescence efficiency ($\Phi$), and fluorescence peak intensity ratio ($R$) (**Figures S8b–e**). Because these four parameters were estimated at the same measurement sites as $\tau_1$ and $\tau_2$, the corresponding distributions are expected to be mutually correlated. This correlation enables further characterization of the $C_B$ and $C_N$ components in terms of these photophysical parameters via the 2D distribution analysis described below.

First, the 2D histograms were constructed by plotting the time constants ($\tau_1$ or $\tau_2$) on the horizontal axis and one of the other parameters (total $\Delta A$, $F$, $\Phi$, or $R$) on the vertical axis (**Figures 5a, c, e, and g**, respectively), with bin sizes of 0.05 ps for $\tau_1$, 0.5 ps for $\tau_2$, $1.0\times10^{-3}$ for total $\Delta A$, 0.3 for $F$, 0.1 for $\Phi$, and 0.1 for $R$. These 2D histograms were fitted with the sum of two 2D Gaussian functions representing the $C_B$ and $C_N$ components, given by:

$$\mathrm{f}(\tau, s) = \sum_{i=\mathrm{C_B},\mathrm{C_N}} A_{2\mathrm{D},i} \exp\left(-\frac{1}{2(1-r_i^2)}\left(\left(\frac{\tau-\tau_{\mathrm{c},i}}{\tau_{\mathrm{w},i}}\right)^2 + \left(\frac{s-s_{\mathrm{c},i}}{s_{\mathrm{w},i}}\right)^2 - \frac{2\cdot r_i\cdot(\tau-\tau_{\mathrm{c},i})\cdot(s-s_{\mathrm{c},i})}{\tau_{\mathrm{w},i}\cdot s_{\mathrm{w},i}}\right)\right), \tag{5}$$

where $i$ labels the $C_B$ or $C_N$ component. The variable $\tau$ is the horizontal-axis coordinate corresponding to either $\tau_1$ or $\tau_2$. The variable $s$ is the vertical-axis coordinate corresponding to one of total $\Delta A$, $F$, $\Phi$, or $R$. $A_{2\mathrm{D},i}$ is the amplitude of the 2D Gaussian distribution for component $i$. $\tau_{\mathrm{c},i}$ and $\tau_{\mathrm{w},i}$ ($s_{\mathrm{c},i}$ and $s_{\mathrm{w},i}$) denote the central position and $1/\sqrt{\mathrm{e}}$ width, respectively, of the $\tau$ ($s$) distribution along the horizontal (vertical) axis for component $i$. $r_i$ indicates the correlation coefficient between $\tau$ and $s$ for component $i$.

Eq. (5) was reformulated for the fitting analysis. The central positions $\tau_{\mathrm{c},i}$ and widths $\tau_{\mathrm{w},i}$ of the $C_B$ and $C_N$ components were fixed at the values in **Table S3**, which were determined from the 2D distribution analysis of the $\tau_1$–$\tau_2$ histogram as described in **Supporting Information Section 8**. Additionally, the relative populations of each component, defined as $P_i$, were constrained to the values in **Table S3**. To implement these constraints, the amplitude $A_{2\mathrm{D},i}$ was rewritten using $P_i$, $\tau_{\mathrm{w},i}$, and $s_{\mathrm{w},i}$, as shown below. First, the integrated value of the 2D Gaussian function for each component, defined as $I_i$, is given by:

$$I_i = 2\pi\cdot A_{2\mathrm{D},i}\cdot\tau_{\mathrm{w},i}\cdot s_{\mathrm{w},i}\cdot\sqrt{1-r_i^2}\,. \tag{6}$$

Here, $I_i$ corresponds to the volume of the 2D distribution for each component, and the total volume of all components is given by $I_{\mathrm{all}} = \sum_i^2 I_i$. Accordingly, the relative population $P_i$ is represented as $P_i = I_i/I_{\mathrm{all}}$. By substituting this relation into eq. (6), $A_{2\mathrm{D},i}$ was rewritten as:

$$A_{2\mathrm{D},i} = \frac{P_i}{\tau_{\mathrm{w},i}\cdot s_{\mathrm{w},i}\cdot\sqrt{1-r_i^2}}\cdot\frac{I_{\mathrm{all}}}{2\pi}\,. \tag{7}$$

By substituting eq. (7) into eq. (5) and replacing the factor $I_{\mathrm{all}}/2\pi$, which is independent of component $i$, with a constant $A_0$, the final fitting function was obtained as follows:

$$\mathrm{f}(\tau, s) = A_0\cdot\sum_{i=\mathrm{C_B},\mathrm{C_N}} \frac{P_i}{\tau_{\mathrm{w},i}\cdot s_{\mathrm{w},i}\cdot\sqrt{1-r_i^2}} \exp\left(-\frac{1}{2(1-r_i^2)}\left(\left(\frac{\tau-\tau_{\mathrm{c},i}}{\tau_{\mathrm{w},i}}\right)^2 + \left(\frac{s-s_{\mathrm{c},i}}{s_{\mathrm{w},i}}\right)^2 - \frac{2\cdot r_i\cdot(\tau-\tau_{\mathrm{c},i})\cdot(s-s_{\mathrm{c},i})}{\tau_{\mathrm{w},i}\cdot s_{\mathrm{w},i}}\right)\right). \tag{8}$$

In this fit, $A_0$ served merely as a scaling factor, and all parameters except $s_{\mathrm{c},i}$, $s_{\mathrm{w},i}$, and $r_i$ were fixed to the previously determined values in **Table S3**. During the fitting analysis, the full width of the distribution along the $s$-axis, given by $2s_{\mathrm{w},i}$, was constrained to be no smaller than the corresponding bin size.

A parameter set of $\tau_1$, $\tau_2$, and $s$, obtained at each measurement site, was binned into the two 2D histograms of $\tau_1$ versus $s$ and $\tau_2$ versus $s$. Consequently, the distribution along the $s$-axis is shared between the two histograms. Therefore, $s_{\mathrm{c},i}$ and $s_{\mathrm{w},i}$, characterizing the $s$ distribution, were constrained to be identical for both the $\tau_1$–$s$ and $\tau_2$–$s$ 2D distributions. Moreover, since both the $C_B$ and $C_N$ components exhibited a strong positive correlation between

$\tau_1$ and $\tau_2$ (**Figure 3c** and **Table S3**), the correlation coefficient between $\tau_1$ and $s$ and that between $\tau_2$ and $s$ are expected to be comparable. To incorporate these considerations, we performed a global fitting analysis of the $\tau_1$–$s$ and $\tau_2$–$s$ 2D distributions, constraining $s_{c,i}$, $s_{w,i}$, and $r_i$ to be shared between them.

In the actual fitting analysis, the fitted results varied substantially depending on the initial values of $s_{c,i}$. Therefore, to determine the optimal initial values, we performed a Monte Carlo simulation in which the fitting was repeated with randomly varied initial values of $s_{c,i}$. As an example, **Figure S10** shows the results obtained for the case where the $s$-axis corresponds to total $\Delta A$. First, the fitting was performed with the initial values of $s_{c,i}$ chosen at random within the range of the $s$-axis for both the $C_B$ and $C_N$ components. Three fitting results obtained using different initial values are shown in **Figures S10a–c**. The results varied depending on the initial values and showed corresponding variations in the $\chi^2$ value. Smaller $\chi^2$ values correspond to better fits. Therefore, we repeated this randomized fitting procedure more than 20,000 times, extracted the trials corresponding to the lowest 1% of $\chi^2$ values, constructed the distribution of the fitted $s_{c,i}$ values as shown in **Figure S10d**, and then estimated the centers of these distributions. By performing the fitting again using the centers as the initial values of $s_{c,i}$, the final results were obtained. The parameter $s$ plotted against the $\tau$-axis was one of total $\Delta A$, $F$, $\Phi$, and $R$, and this Monte Carlo-based procedure was performed for each choice of $s$.

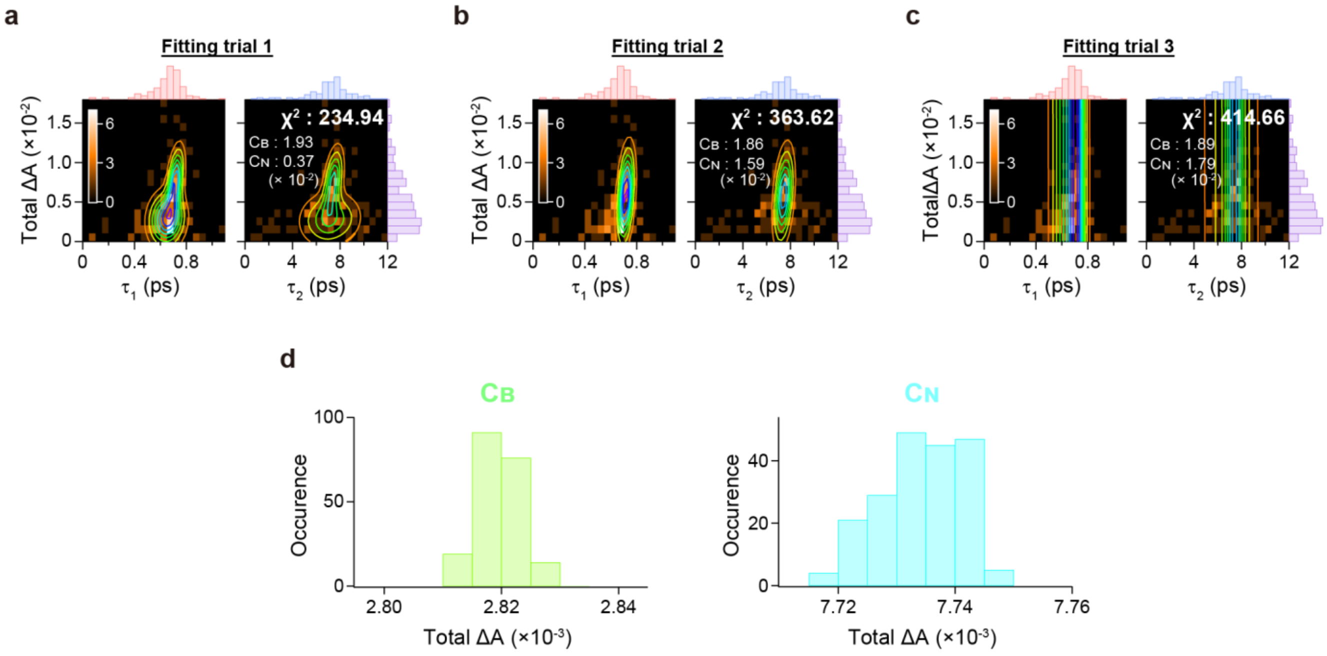


**Figure S10.** 2D distribution analysis using two-component 2D Gaussian fits with Monte Carlo simulation. (a–c) Three representative results obtained from a global fit using two-component 2D Gaussian functions for the 2D histograms of $\tau_1$ versus total $\Delta A$ and $\tau_2$ versus total $\Delta A$, in which the initial total $\Delta A$ values were randomly assigned to the $C_B$ and $C_N$ components. The fitted 2D distributions are shown as contours in each panel. The initial total $\Delta A$ values and $\chi^2$ values for these fits are annotated in each panel. Projections of the distributions onto the $\tau_1$, $\tau_2$, and total $\Delta A$ axes are shown at the top of the left and right panels and at the right of the right panel, respectively. (d) Distributions of total $\Delta A$ estimated for the $C_B$ (left) and $C_N$ (right) components from the fits, constructed from the trials corresponding to the lowest 1% of $\chi^2$ values out of more than 20,000 trials.

**(ii) Calculation of *p*-values from statistical tests**

As described above, the central position $s_{c,i}$ and $1/\sqrt{e}$ width $s_{w,i}$ of the distributions along the $s$-axis were determined for the $C_B$ and $C_N$ components. To assess whether the *s* distributions for the $C_B$ and $C_N$ components exhibit a statistically significant difference, Welch's *t*-test was performed. The *t*-statistic for evaluating the difference in the central positions between components *i* and *j* is given by:

$$t = \frac{s_{c,i} - s_{c,j}}{\sqrt{\frac{u_i^2}{n_i} + \frac{u_j^2}{n_j}}}\,, \tag{9}$$

where *i* and *j* label the $C_B$ or $C_N$ components. $s_{c,i}$ ($s_{c,j}$) is the central position along the *s*-axis for component *i* (*j*), estimated from the 2D distribution fit. $n_i$ ($n_j$) is the sample size of each *s* distribution for component *i* (*j*), which was calculated by multiplying the relative population $P_i$ ($P_j$) by the total number of data points included in the entire 2D distribution. $u_i^2$ ($u_j^2$) is the unbiased variance for component *i* (*j*), which was obtained from the width $s_{w,i}$ ($s_{w,j}$) as follows:

$$u_i^2 = \left(s_{w,i}\frac{n_i}{n_i - 1}\right)^2 . \tag{10}$$

Additionally, the degrees of freedom (DOF) were calculated using the equation below:

$$\mathrm{DOF} = \frac{\left(\frac{u_i^2}{n_i} + \frac{u_j^2}{n_j}\right)^2}{\frac{\left(\frac{u_i^2}{n_i}\right)^2}{n_i - 1} + \frac{\left(\frac{u_j^2}{n_j}\right)^2}{n_j - 1}} . \tag{11}$$

From the *t*-statistic and the DOF, the *p*-value for the difference between the *s* distributions for components *i* and *j* was calculated. A small *p*-value indicates statistical significance.

Moreover, we tested the statistical significance of the correlation coefficient $r_i$. Assuming that the population correlation coefficient is $r = 0$, the *t*-statistic for the sample correlation coefficient $r_i$, denoted as $t_i$, is given by:

$$t_i = r_i\sqrt{\frac{n_i - 2}{1 - r_i^2}} . \tag{12}$$

Considering that $t_i$ follows a *t* distribution with $n_i - 2$ degrees of freedom, the *p*-value for $r_i$ was calculated. A small *p*-value indicates that $r_i$ is significantly different from $r = 0$.

**Supporting Information Section 11: Fluorescence lifetime measurements**

To estimate the excited-state lifetime, time-resolved fluorescence measurements of individual Zn-HM aggregates were performed with detection at >750 nm, using an avalanche photodiode (APD) with low jitter (PDM, Micro Photon Devices) connected to a time-correlated single-photon counting (TCSPC) module (PicoHarp300, PicoQuant). The sample was excited at ~695 nm (**Figure 2d**, blue) with a fluence of $3.9 \times 10^{14}$ photons pulse$^{-1}$ cm$^{-2}$. As shown in **Figure S11**, the fluorescence decay profiles (red) obtained at three distinct sites in different aggregates were nearly identical to the IRF (black). Fitting each decay with a single-exponential function convolved with the IRF yielded a fluorescence lifetime $\tau_F$ at each site (blue). The mean $\tau_F$ across the three sites was 26 ps with a standard deviation of 5 ps. These results suggest that the excited state of the Zn-HM aggregates relaxes to the ground state or transitions to a non-emissive dark state within the temporal resolution corresponding to the FWHM of the IRF (~40 ps). The excitation dynamics leading to fluorescence emission are discussed in the main text and **Supporting Information Section 12**.

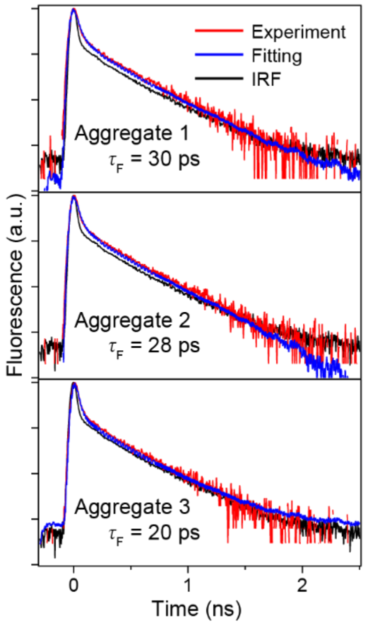


**Figure S11.** Fluorescence lifetime measurements of individual Zn-HM aggregates. Fluorescence decay profiles (red) acquired at >750 nm from individual Zn-HM aggregates at different sites are shown in each panel. The same IRF profile (black), acquired in this wavelength range, is shown in all panels. The vertical axes are displayed on a log scale. The fluorescence lifetime $\tau_F$, estimated from fits using a single-exponential function convolved with the IRF (blue), is annotated in each panel.

**Supporting Information Section 12: Heterogeneous aggregate model incorporating exciton domains in J-aggregates**

The photophysical properties of the $C_B$ and $C_N$ components obtained from the 2D distribution analysis (**Figure 5**) can be interpreted in terms of the heterogeneous aggregate models schematically illustrated in **Figure S12**. The aggregates form layered structures composed of a large number of Zn-HM molecules (oriented gray bars), in which exciton domains (yellow circles) are generated by the mixing of multiple excited states mediated by intermolecular interactions. The domain size is restricted to a finite scale because the excitonic coherence length is limited by microscopic disorder arising from structural distortions and fluctuations.[5-6] Consequently, multiple exciton domains are considered to be spatially and energetically dispersed throughout the aggregates. When one of these exciton domains absorbs light (red arrows), excitation energy migrates and diffuses among the exciton domains (black arrows) and is finally emitted as fluorescence or dissipated as thermal energy (blue arrows). By confining the measurement region to approximately the diffraction limit (black outline), the observed heterogeneity can be spatially restricted, thereby enabling us to discuss the excitation dynamics in relation to the microscopic disorder within the aggregates.

The $C_N$ component, exhibiting a narrow $\tau$ distribution, was identified as originating from structural regions with relatively homogeneous molecular packing. Within such regions, the following two types of local configurations are likely distributed. Some regions contain exciton domains of nearly uniform size (**Figure S12a**), and others contain a mixture of large and small exciton domains (**Figure S12b**). In the former case, molecular clusters that form exciton domains of similar size are considered to have comparable energy levels, suggesting that the absorbed excitation energy can diffuse within the measurement region. Therefore, light absorption and subsequent fluorescence emission and thermal dissipation are governed by exciton domains of similar size, eventually returning the system to the ground state. In contrast, in the latter case, the excitation energy is trapped by a large exciton domain with a longer coherence length, i.e., the molecular cluster with the lowest energy level. Accordingly, the fluorescence spectrum and fluorescence efficiency primarily reflect the properties of exciton domains with longer coherence lengths. Conversely, the TA signals predominantly reflect the properties of smaller exciton domains with shorter coherence lengths.

When large exciton domains are responsible for both light absorption and fluorescence emission (**Figure S12c**), absorption and fluorescence measurements should exhibit behavior reflecting a long coherence length. Specifically, both the non-radiative decay time constant $\tau$ and the fluorescence efficiency $\Phi$ are predicted to increase. In this study, time constants exceeding the 12-ps time window were observed at ~14% of the measurement sites. For these sites, the fluorescence efficiency $\Phi$ was 2.0 with a standard deviation of 1.0, and the fluorescence peak intensity ratio $R$ was 1.6 with a standard deviation of 0.5. Accordingly, these sites exhibited larger $\tau$, larger $\Phi$, and similar (or slightly larger) $R$ values compared to those of the $C_N$ component, suggesting that both light absorption and fluorescence emission occur in exciton domains with longer coherence lengths. Thus, the >12 ps component is likely attributable to regions predominantly containing large exciton domains.

In the regions exhibiting the $C_B$ component, which shows a broad $\tau$ distribution, the structural heterogeneity within the measurement spot is larger than that in the regions exhibiting the $C_N$ component, thereby suggesting a mixture of exciton domains of various sizes (**Figure S12d**). Accordingly, after light absorption, the excitation energy is trapped at exciton domains with low energy levels and long coherence lengths, from which fluorescence is emitted. In this situation, even if the measurement spot is confined to the diffraction limit, the obtained results reflect a subensemble average, obscuring the correlations between parameters.

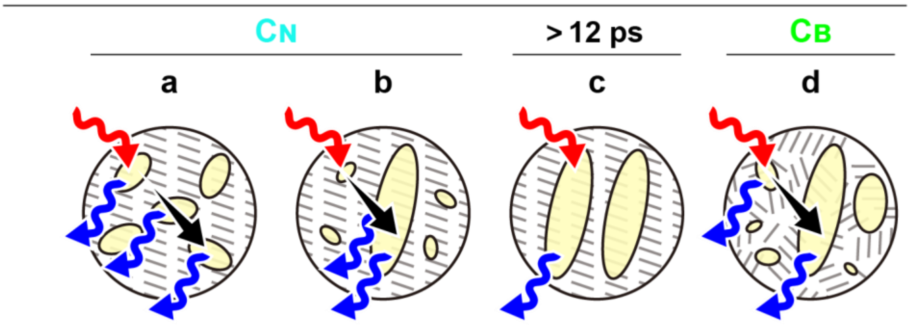


| | a | b | c | d |
|---|---|---|---|---|
| $\tau$ | L | S | VL | M |
| $\Delta A$ | L | S | VS | S |
| $F$ | L | S | VS | S |
| $\Phi$ | S | L | VL | M |
| $R$ | S | L | L | M |

**Figure S12.** Heterogeneous aggregate model incorporating exciton domains in J-aggregates. Microscopic aggregate architectures assigned to the $C_N$ (a, b), $\tau$ > 12 ps (c), and $C_B$ (d) components, observed within the diffraction-limited spot (black outlines), are shown. Zn-HM molecules (oriented gray bars) self-assemble into J-aggregates, in which exciton domains formed by intermolecular interactions (yellow circles) are distributed. Light absorption (red arrows), excitation energy diffusion (black arrows), and fluorescence emission or thermal dissipation (blue arrows) are mediated by these exciton domains. The relative magnitudes of photophysical properties such as $\tau$, total $\Delta A$, $F$, $\Phi$, and $R$, as predicted by each model, are listed at the bottom. VL: very large, L: large, M: medium, S: small, VS: very small, where M corresponds to the mean magnitude, i.e., the average across (a) and (b), of each property for $C_N$. For $C_B$, these magnitudes are broadly distributed and are therefore indicated as subensemble averages.
